# RAP1 regulates TIP60 function during fate transition between 2 cell-like and pluripotent states

**DOI:** 10.1101/2021.11.02.467017

**Authors:** Raymond Mario Barry, Olivia Sacco, Amel Mameri, Martin Stojaspal, William Kartsonis, Pooja Shah, Pablo De Ioannes, Ctirad Hofr, Jacques Côté, Agnel Sfeir

## Abstract

In mammals, the conserved telomere binding protein RAP1 serves a diverse set of non-telomeric functions including activation of the NF-kB signaling pathway, maintenance of metabolic function *in vivo,* and transcriptional regulation. Here, we uncover the mechanism by which RAP1 modulates gene expression. Using a separation-of-function allele, we show that RAP1 transcriptional regulation is independent of TRF2-mediated binding to telomeres and does not involve direct binding to genomic loci. Instead, RAP1 interacts with the TIP60/p400 complex and modulates its histone acetyltransferase activity. Notably, we show that deletion of RAP1 in mouse embryonic stem cells increases the fraction of 2-cell-like cells. Specifically, RAP1 enhances the repressive activity of Tip60/p400 across a subset of 2-cell-stage genes, including *Zscan4* and the endogenous retrovirus MERVL. Preferential upregulation of genes proximal to MERVL elements in Rap1 deficient settings implicate these endogenous retroviral elements in the de-repression of proximal genes. Altogether, our study reveals an unprecedented link between RAP1 and TIP60/p400 complex in the regulation of totipotency.

## Introduction

Rap1 (Repressor/ Activator Protein 1) was first identified in *Saccharomyces cerevisiae* (*S. cerevisiae*) as a transcription factor that silences the alternate silent mating-type loci and maintains high transcriptional activity at genes encoding ribosomal proteins and glycolytic enzymes (Huet et al. 1985; Huet and Sentenac 1987; Shore and Nasmyth 1987; Shore et al. 1987; Vignais et al. 1987; Chambers et al. 1989). Subsequent studies found yeast Rap1 to be a primary telomere DNA binding protein that maintains proper telomere length, ensures subtelomere silencing, and inhibits telomere end-end fusions (Berman et al. 1986; Buchman et al. 1988a; Buchman et al. 1988b; Longtine et al. 1989; Lustig et al. 1990; Kyrion et al. 1993; Pardo and Marcand 2005). Upon senescence of budding yeast, Rap1 is evicted from telomeres and relocalizes to promoter targets, including histone-encoding genes (Platt et al. 2013; Song et al. 2020). In *Schizosaccharomyces pombe* (*S. pombe*), Rap1 performs a similar protective function at telomeres, albeit, it relies on a protein-protein interaction with Taz1 to be recruited to chromosome ends (Kanoh and Ishikawa 2001).

Mammalian RAP1, encoded by *TERF2IP*, is a component of shelterin, a six-subunit protein complex that is additionally composed of TRF1, TRF2, TIN2, POT1, and TPP1. Shelterin coats mammalian telomere DNA and prevents the activation of DNA damage signaling and repair at chromosome ends (de Lange 2005; Lazzerini-Denchi and Sfeir 2016). Whereas budding yeast Rap1 binds telomeric DNA directly using two tandem Myb domains, mammalian RAP1 comprises a single Myb domain and relies on a stable interaction with TRF2 to be recruited to TTAGGG-bearing telomere DNA (Li et al. 2000; Hanaoka et al. 2001). RAP1 function at telomeres diverged from yeast as revealed by genetic studies showing that deletion of mammalian RAP1 is largely dispensable for telomere end-protection. Whereas loss of all other shelterin components is incompatible with life, Rap1 knockout (*Rap1^-/-^*) mice are alive, fertile and display no premature aging phenotypes (Karlseder et al. 2003; Chiang et al. 2004; Celli and de Lange 2005; Hockemeyer et al. 2006; Kibe et al. 2010; Sfeir et al. 2010). Loss of function analysis indicated that RAP1 is dispensable for chromosome-end protection. However, in certain experimental settings, it was found to act with TRF2 to suppress NHEJ mediated telomere fusions (Sarthy et al. 2009; Kabir et al. 2014; Lototska et al. 2020). Furthermore, in the absence of Ku70/80, telomere sister-chromatic exchanges are induced upon Rap1 depletion, implicating a redundant role for RAP1 in suppression of telomere recombination (Sfeir et al. 2010; Rai et al. 2016).

Analysis of *Rap1^-/-^* mice *in vivo* and *ex vivo* shed light into the function of mammalian Rap1 in regulating gene expression (Martinez et al. 2010; Teo et al. 2010; Martinez et al. 2013; Yeung et al. 2013). At the organismal level, Rap1 deficiency leads to glucose intolerance, dyslipidemia, liver steatosis, and excess fat accumulation, ultimately manifesting as late-onset obesity. *Rap1^-/-^* mice exhibit altered gene expression in a subset of tissues and prior to the onset of obesity. Furthermore, mouse embryonic fibroblasts (MEFs) from *Rap1^-/-^* embryos display dysregulated gene expression, a phenotype rescued by exogenous expression of Rap1, indicating that Rap1 regulates gene expression in a cell-autonomous manner (Martinez et al. 2013; Yeung et al. 2013). These observations point to an extratelomeric function of RAP1 in regulating gene expression; however, the mechanistic basis of this regulation remains elusive.

Regulation of gene expression is orchestrated by a complex interplay of transcription factors and chromatin regulators, including chromatin remodelers, histone chaperones, and epigenetic modifiers that write/erase histone post-translational modifications. Histone acetylation is one such modification that regulates the structure of chromatin and typically leads to activation of gene expression (Workman and Abmayr 2014). Histone acetyltransferase (HAT) complexes tend to be large and multimeric, such as the highly conserved TIP60/p400 complex, also known as NuA4. TIP60/p400 is composed of >18 subunits including the lysine acetyltransferase (KAT) catalytic subunit Tip60 (*KAT5*), two scaffold proteins p400 (*Ep400*) and Trrap, Epc1/2, Dmap1 and Ing3 (Doyon and Cote 2004). TIP60/p400 plays a role in a diverse set of biological processes including transcription, cell proliferation, and DNA repair by homologous recombination (Steunou et al. 2014; Jacquet et al. 2016; Sheikh and Akhtar 2019). TIP60/p400 regulates transcription through acetylation of histones H4 and H2A catalyzed by the TIP60 subunit, but can also modulate gene expression by incorporation of the histone variant H2A.Z using the ATP-dependent chromatin remodeling activity of Ep400 (Allard et al. 1999; Pradhan et al. 2016). TIP60/p400 is also necessary for proper maintenance and renewal of stem cells (Fazzio et al. 2008; Acharya et al. 2017). Specifically, this HAT complex was recently found to suppress emergence of 2-cell-like (2C-like) cells, a small population of cells in mouse embryonic stem cell (mESC) cultures that share characteristics with two-cell stage embryos including expression of endogenous retroviral MERVL elements and enhanced totipotent potential (Rodriguez-Terrones et al. 2018). Paradoxically, TIP60/p400 increases totipotency by repressing a subset of 2C genes, however, the underlying basis of this noncanonical TIP60 function in suppressing transcription remains elusive.

Here, we uncover the mechanism by which endogenous RAP1 maintains native gene expression. We show that RAP1 transcriptional function in mammalian cells is independent of its telomere association and does not involve binding to DNA. Instead, our data reveal that non-telomeric RAP1 interacts with the TIP60/p400 complex by binding cooperatively to TIP60 and Epc1 and enhances TIP60 acetylation activity. Notably, loss of RAP1 in mESCs triggers a 2C-like state and modulates the repressive activity of TIP60/p400 on a subset of 2C-stage genes, especially ones proximal to the endogenous retroviral MERVL elements. In summary, our data uncover an unprecedented link between a conserved telomere binding protein and the TIP60/p400 complex in the regulation of totipotency.

## Results

### Endogenous Rap1 maintains native transcription independent of telomeres

It has been proposed that the function of mammalian RAP1 in gene regulation is similar to budding yeast Rap1 and driven by its association with promoter elements. Consistent with this hypothesis, a study hinted at direct binding of mammalian RAP1 to DNA in a sequence non-specific manner (Arat and Griffith 2012), and ChIP-seq analysis highlighted several genomic loci that are bound by Rap1 (Martinez et al. 2010; Yang et al. 2011). To gain better insight into the mechanism by which Rap1 regulates transcription, we employed CRISPR/Cas9 to generate mice carrying *Rap1^I312R^* (Figure 1A), an allele that is incapable of binding TRF2 (Chen et al. 2011). Fertilized oocytes were injected with mRNA encoding Cas9 nuclease, a single-guide RNA (sgRNA) targeting exon 3, and a single-stranded DNA oligo donor (ssODN) coding for *Rap1^I312R^*. Zygotes were implanted into surrogate female mice, and proper targeting was confirmed by genotyping PCR and Sanger sequencing (Figure 1B-C).

**Figure 1.**
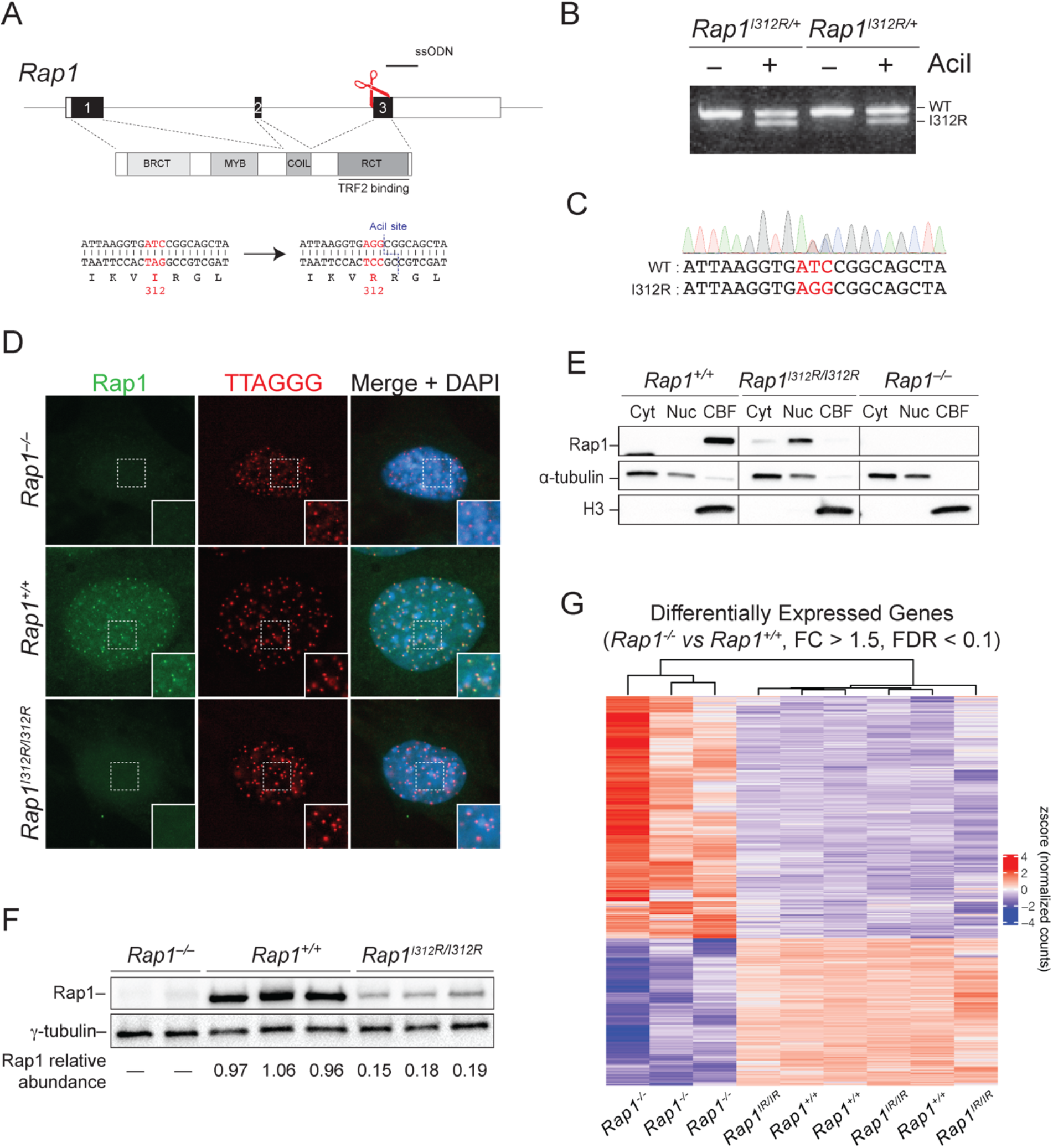
Rap1 maintains gene transcription independent of telomeres. A) Schematic representation of CRISPR/Cas9 gene editing strategy to substitute Rap1 isoleucine 312 to arginine. Successful targeting creates an AciI restriction site to be used during genotyping. Single-stranded oligo donor (ssODN, black line) single-guide RNA (sgRNA) cut site (red scissors) are indicated. B) PCR genotyping of tail-tip DNA from two *Rap1^I312R/+^* heterozygous mice following gene targeting. C) Example Sanger sequencing of *Rap1^I312R/+^* heterozygous mouse. D) Representative image of IF-FISH in *Rap1^+/+^*, *Rap1^-/-^*, and *Rap1^I312R/I312R^* (labelled as *Rap1^IR/IR^*) MEFs for Rap1 (green) and telomeres (red) using Rap1 antibody and TTAGGG PNA probe, respectively. DAPI (blue) is used as counterstain. E) Rap1 Western blot from *Rap1^-/-^*, *Rap1^+/+^*, and *Rap1^I312R/I312R^* MEFs following subcellular fractionation into cytoplasmic (Cyt), nucleoplasmic (Nuc), and chromatin-bound (CBF) fractions. *α*-tubulin and histone H3 are loading controls for Cyt and CBF, respectively. Blot is representative of n = 2 biological replicates. F) Immunoblot for Rap1 from whole cell lysates obtained from *Rap1^-/-^* (n = 2), *Rap1^+/+^* (n = 3), and *Rap1^I312R/I312R^* (n = 3) MEFs. Rap1 relative abundance was determined by normalizing to *γ*-tubulin. G) Hierarchically clustered heatmap representing RNA-seq data for differentially expressed genes (DEGs) between *Rap1^+/+^* and *Rap1^-/-^* MEFs (FC > 1.5; FDR < 0.1; n = 3 biological replicates per genotype).

To confirm the release of RAP1 from telomeres in *Rap1^I312R/I312R^* (also labeled *Rap1^IR/IR^*) mouse embryonic fibroblasts (MEFs), we performed indirect immunofluorescence coupled with fluorescence *in situ* hybridization (IF-FISH). Consistent with previous reports (Chen et al. 2011; Yeung et al. 2013), Rap1 exhibited a diffuse nucleoplasmic staining pattern in *Rap1^I312R/I312R^* MEFs and contrasted with the punctate staining that co-localized with telomeres in *Rap1^+/+^* cells (Figure 1D). We then fractionated lysates into cytosolic, nucleoplasmic, and chromatin-bound fractions and showed that whereas wild type RAP1 was strongly bound to chromatin, Rap1^I312R^ lost its chromatin association and as distributed between the nucleoplasm and cytoplasm (Figure 1E). Furthermore, Western blot analysis revealed that RAP1 protein abundance was reduced ∼ 5-fold in *Rap1^I312R/I312R^* MEFs relative to wild type cells (Figure 1F). Similarly, overall Rap1 levels were reduced when examined in different tissues harvested from *Rap1^I312R/I312R^* mice (Figure S1A-B). This observation is in accordance with previous reports indicating that binding to TRF2 stabilizes RAP1 protein levels (Celli and de Lange 2005; Sfeir et al. 2010). Along these lines, treatment with the proteasome inhibitor, MG132 (5 µM for 11 hours) increased RAP1 abundance by two- and six-fold in *Rap1^+/+^* and *Rap1^I312R/I312R^* MEFs, respectively (Figure S1C-D). In addition, co-immunoprecipitation (CoIP) experiments in HEK293T cells showed that Rap1 immunoprecipitated with K48Ub, providing further evidence for its K48-linked ubiquitinylation (Figure S1E). Altogether, our results show that at telomeres, mammalian Rap1 is stabilized by its interaction with TRF2, while the extratelomeric RAP1 pool is tightly regulated by proteasomal degradation.

We then performed RNA sequencing (RNA-Seq) on *Rap1^-/-^*, *Rap1^+/+^* and *Rap1^I312R/I312R^* primary MEFs. Principal component analysis (PCA) of transcriptomes showed close clustering of the *Rap1^+/+^* and *Rap1^I312R/I312R^* samples apart from *Rap1^-/-^* samples (Figure S1F). Differential gene expression analysis using DESeq2 identified 654 differentially expressed genes in *Rap1^-/-^* vs. *Rap1^+/+^* MEFs (FC > 1.5, FDR < 0.1; Supplementary Table S1). These genes were involved in various pathways related to cell-cell signaling and developmental processes (Figure S1G; Supplementary Table S2). Interestingly, gene expression (653 of 654, 99.8%) was largely maintained in *Rap1^I312R/I312R^* MEFs relative to wild type cells (Figure 1G). Taken together, our data suggest that despite the overall reduction in Rap1 levels in *Rap1^I312R/I312R^* cells, a small pool of extra-telomeric RAP1 is sufficient to fully maintain gene expression control. Furthermore, we establish that the function of mammalian RAP1 in gene expression is completely independent of TRF2 binding and telomere localization.

### Rap1^I312R^ does not directly associate with chromatin

In budding yeast, extra-telomeric RAP1 binds directly to interstitial genomic sites including several gene promoters (Bram and Kornberg 1985; Huet et al. 1985; Shore and Nasmyth 1987; Vignais et al. 1987; Chambers et al. 1989; Platt et al. 2013; Song et al. 2020). Similarly, in human and mouse cells, wild type Rap1 was detected at many sites throughout the genome (Martinez et al. 2010; Yang et al. 2011; Martinez et al. 2016) with a notable overlap with TRF2 binding sites. Given that Rap1^I312R^ fully maintains transcriptional regulation, we sought to employ this separation-of-function allele to identify TRF2-independent RAP1 binding sites and, thus, more relevant to its role in gene expression. We transduced SV40LT-immortalized *Rap1^-/-^* MEFs with HA-Rap1 and HA- Rap1-I312R and noted that both alleles accumulated at similar levels when exogenously expressed (Figure S2A). As predicted, HA-Rap1 localized to telomeres whereas HA-Rap- I312R was largely nucleoplasmic (Figure S2B). Chromatin immunoprecipitation (ChIP) using anti-HA antibody followed by dot blot analysis showed significant association of telomeric DNA with HA-RAP1 but not with HA-RAP1-I312R (Figure 2A). In addition, analysis by high-throughput sequencing revealed a 100-fold enrichment for reads containing telomere repeats in cells expressing wild type Rap1 (Figure 2B). We performed peak calling using MACS2 (q < 0.05) and identified 109 wild type Rap1 binding sites: 39 subtelomere sites and 70 other loci scattered throughout the genome (Figure 2C, representative peaks Figure S2C-D; Supplementary Table S3). This analysis is consistent with previous observations that RAP1 localizes to discrete sites throughout the genome (Martinez et al. 2010; Yang et al. 2011; Martinez et al. 2016). Notably, using broad as well as narrow peak calling algorithms, we did not identify any genomic binding sites for RAP1^I312R^ (Figure 2C). Specifically, RAP1^I312R^ showed no binding near telomeres (Figure 2D) nor at interstitial peaks bound by wild type Rap1 (Figure 2E).

**Figure 2.**
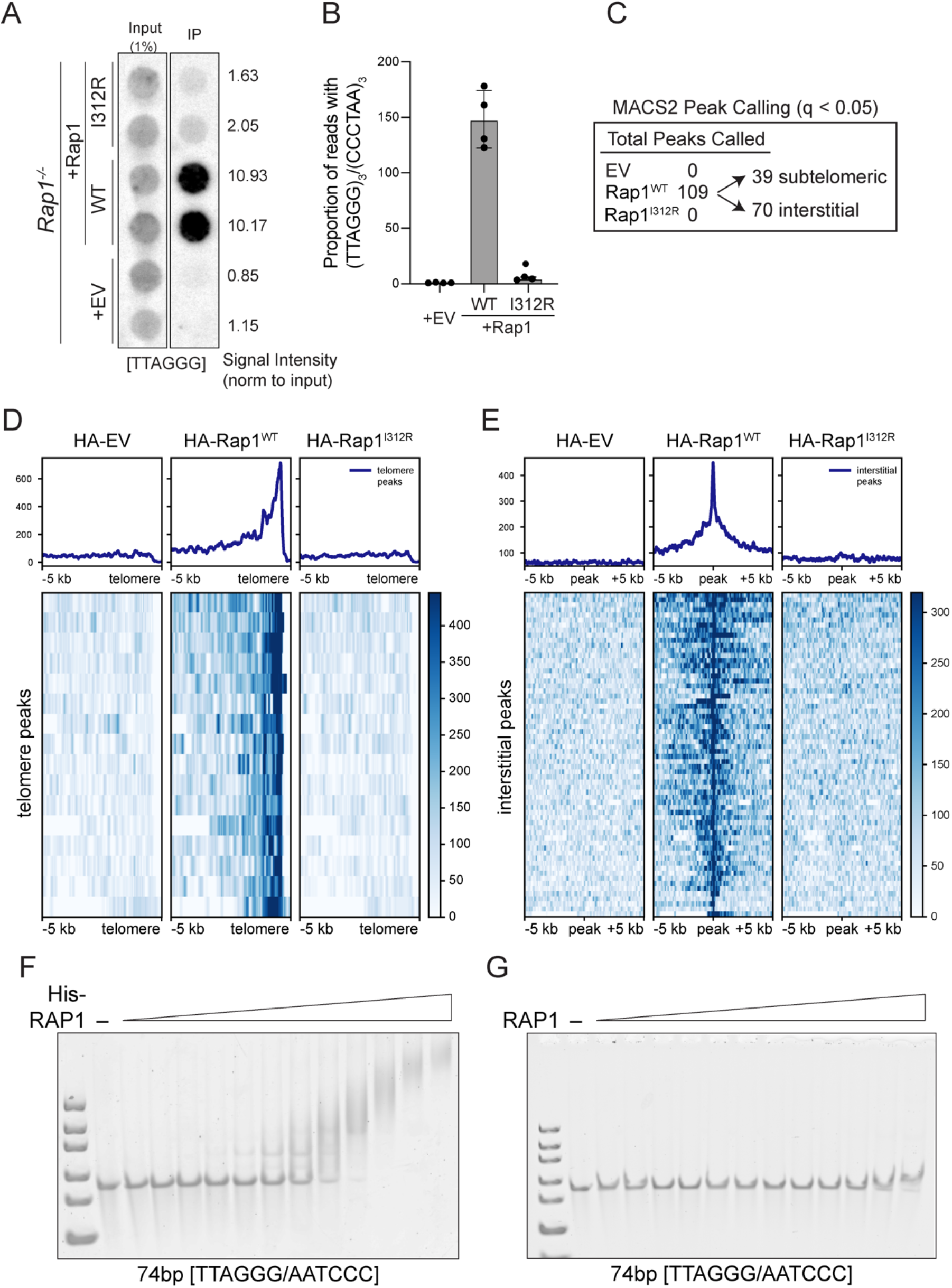
Rap1-I312R does not bind to genomic loci or DNA. A) Dot blot for telomere repeats following ChIP using anti-HA antibody in cells expressing HA-tagged Rap1-WT, Rap1-I312R, or empty vector (EV) control (n = 2 biological replicates). Signal intensity is determined by normalizing to input. B) High-throughput sequencing of ChIP samples as in *A*. Telomere binding was determined by calculating the proportion of reads with at least three telomere (TTAGGG)_3_/(CCCTAA)_3_ repeats. Data are the mean ± standard deviation of n = 4 biological replicates. C) Summary of ChIP-seq peak calling analysis which identified 109 peaks in HA-Rap1-WT samples (MACS2; q <0.05). Peaks were classified as subtelomeric if localized within 500kb from the telomere. D) Heatmap representing ChIP-seq profiles of HA-Rap1-WT peaks at chromosome ends. E) Heatmap representing ChIP-seq profiles of HA-Rap1-WT interstitial peaks. F) Electrophoretic mobility shift assay (EMSA) using His-tagged Rap1 (0 to 15 !M) and a 74bp TTAGGG/AATCCC double-stranded DNA (100 nM). G) EMSA using non-His-tagged-Rap1 (0 to 15 !M) and a 74bp TTAGGG/AATCCC double-stranded DNA (100 nM).

Having failed to detect RAP1^I312R^ binding to DNA by ChIP, we then turned to *in vitro* assays to directly assess the binding of Rap1 to DNA. We performed electrophoretic mobility shift assays (EMSAs) using purified protein and initially found that His-tagged RAP1 binds to DNA with no sequence specificity and is also capable of associating with purified nucleosomes (Figure 2F, S2E-F). Importantly, upon removal of the His-tag used for purification purposes, the binding of RAP1 to free and nucleosome-bound DNA was abrogated (Figure 2G; Figure S2G-H). The confounding impact of histidine tags on DNA-binding activity of purified proteins has been reported in the literature (Paul et al. 2020) and could well explain earlier studies that showed nonspecific binding of human RAP1 to DNA *in vitro* (Arat and Griffith 2012). Taken together, our data confirm that unlike budding yeast Rap1, the mammalian counterpart is incapable of binding DNA directly, and therefore, the mechanism by which RAP1 regulates gene expression in higher eukaryotes must have diverged from budding yeast.

### Proximity-based biotinylation reveals extratelomeric Rap1 interactors

We next used proximity-based biotinylation to identify RAP1 protein interactions and provide insight to the potential mechanisms by which extratelomeric Rap1 regulates gene expression (Roux et al. 2012). To that end, we transduced SV40LT-immortalized *Rap1^-/-^* MEFs with BioID-tagged RAP1 and RAP1-I312R and established independent clonally derived cells that expressed similar levels of the RAP1 variants. We reasoned that exploring the RAP1 proteome in the context of RAP1-I312R will enable the identification of extra-telomeric RAP1 interactors relevant to its function in gene regulation. As expected, BioID-RAP1 colocalized with telomere TTAGGG (Figure 3A) and interacted with TRF2 (Figure 3B), whereas BioID-RAP1-I312R expressed diffusely throughout the nucleus (Figure S3A) and failed to associate with TRF2 (Figure 3B).

**Figure 3.**
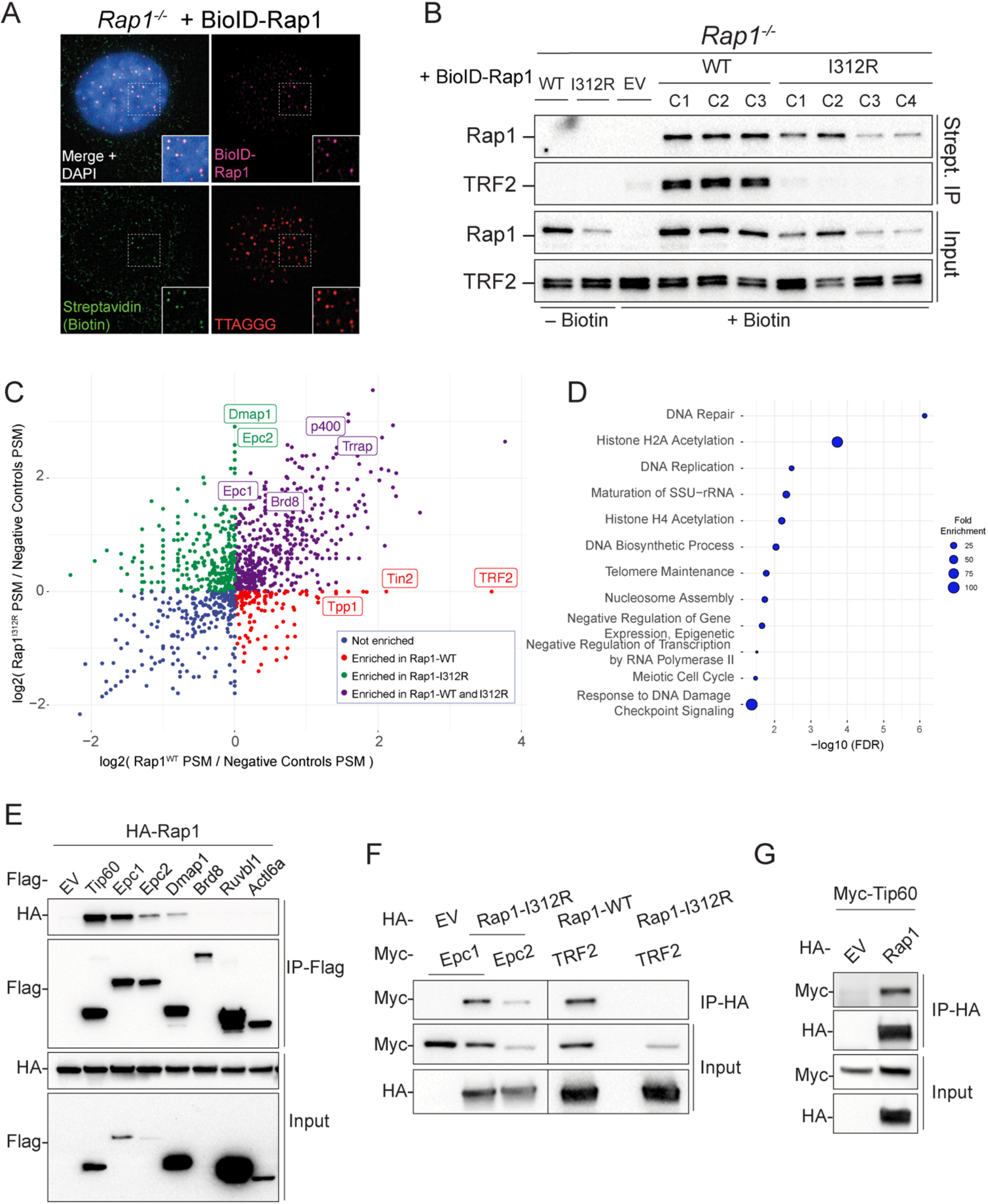
Proximity-dependent biotinylation reveals extratelomeric Rap1 binding partners. A) Representative image of IF-FISH using streptavidin antibody in *Rap1^-/-^* MEFs expressing BioID-Rap1-WT stained for BioID-Rap1 (magenta), biotin (green), and telomeres (red) using anti-Flag antibody, streptavidin, and TTAGGG PNA probe, respectively. DAPI (blue) is used as counterstain. B) Immunoblot for Rap1 and TRF2 following streptavidin pull-down in *Rap1^-/-^* MEFs expressing BioID-Rap1-WT (n = 3 biological replicates; clones C1, C2, C3), BioID-Rap1-I312R (n = 4 biological replicates; clones C1, C2, C3, C4), and BioID alone (EV). Where indicated, cells were treated with biotin (50 μM) for 20 hours prior to harvest. C). Scatter plot representing log_2_ fold change in peptide-spectrum match (PSM) of proteins identified by BioID-Rap1-WT versus BioID- Rap1-I312R. Fold change values were calculated relative to no biotin and BioID alone controls. Each dot represents a unique protein. Known Tip60/p400 (green, purple) and shelterin (red) complex members are indicated. D) Graphical representation of top 10 biological processes overrepresented in BioID-Rap1-I312R streptavidin pull-down (FC > 2) using PANTHER Classification System statistical overrepresentation test. E) CoIP of Myc-tagged Tip60/p400 subunits (Tip60, Epc1, Epc2, Dmap1, Brd8, Ruvbl1, Actl6a) and HA-Rap1 following co-transfection in HEK293T cells. F) Reciprocal CoIP of HA-Rap1- I312R and Myc-tagged Epc1 or Epc2. CoIP of HA-Rap1 or HA-Rap1-I312R with TRF2 is used as a control. G) Reciprocal CoIP of HA-Rap1 and Myc-Tip60.

We then performed mass spectrometry analysis following streptavidin pull-down of nuclear extracts (silver stain and streptavidin-HRP, Figure S3B; Supplementary Table S4) and detected 189 and 227 interactors of RAP1 and RAP1-I312R, respectively, that were enriched at least 2-fold (peptide-spectrum match for protein abundance) over the empty vector (BioID) control (Supplementary Table S5). As expected, proteins exclusively associating with wild type RAP1 included shelterin subunits TRF2, TPP1, and TIN2, (Figure 3C; Figure S3D). Notably, among factors enriched in the context of Rap1-I312R, we identified six members of the TIP60/p400 histone acetyltransferase complex, EPC1 and its paralog EPC2, DMAP1, p400, TRRAP, and BRD8 (Figure 3C; Figure S3C). Furthermore, statistical overrepresentation test of biological processes using PANTHER Classification System revealed statistical significance for Histone H2A Acetylation, Histone H4 Acetylation, and DNA Repair in RAP1-I312R (Figure 3D; Supplementary Table S6).

To validate the mass spectrometry data, we performed CoIP of Flag- and HA-tagged proteins co-expressed in HEK293T and demonstrated stable interaction between RAP1 and TIP60, EPC1, EPC2, and DMAP1 (Figure 3E-G). Importantly, the interaction between RAP1 and EPC1 was not diminished upon treatment with DNase (Figure S3D), and was evident when using purified recombinant proteins (GST-EPC1 and His-RAP1) (Figure S3E), suggesting that the two proteins interact independent of DNA binding. In conclusion, our results reveal that extra-telomeric RAP1 is a binding partner of the TIP60/p400 complex.

### Rap1 associates with Epc1 and Tip60, enhancing histone acetyltransferase (HAT) activity

The TIP60/p400 histone acetyltransferase is a multimeric complex composed of >18 subunits (Doyon and Cote 2004). TRRAP and p400 serve as scaffolds for the assembly of the complex whereas TIP60, a lysine acetyltransferase (KAT), is the catalytic subunit responsible for acetylation of histones H2A, H4, H2A.Z and other nonhistone proteins, including Tp53 (Steunou et al. 2014). Epc1 comprises four conserved domains (Figure 4A). The N-terminal EPcA domain assembles the core 4-subunit catalytic module that includes Tip60 (a.k.a. piccolo NuA4 or picNuA4), while the central EPcB domain binds p400 to effectively bridge the rest of the TIP60/p400 complex (Boudreault et al. 2003; Doyon et al. 2004; Selleck et al. 2005; Auger et al. 2008; Chittuluru et al. 2011; Xu et al. 2016; Setiaputra et al. 2018). Using deletion mutants of the Epc1 subunit, we found that the EPcA domain is necessary and sufficient to interact with Rap1 (Figure 4A-C), placing the telomere binding protein at the Epc1/TIP60 interaction interface. Through a reciprocal approach, we identified the C-terminal RCT domain of Rap1 to be necessary for binding to EPC1 (Figure 4D). The RAP1 RCT domain also binds TRF2, suggesting that binding of RAP1 to TIP60/p400 and TRF2 is mutually exclusive. Consistent with this notion, we found no evidence for TRF2 binding to EPC1/RAP1, in contrast TRF2 reduces the EPC1-RAP1 interaction (Figure S4A). Taken together, these data implicate RAP1 in two separate complexes, being part of shelterin at telomeres and associating with TIP60/p400 during transcriptional regulation.

**Figure 4.**
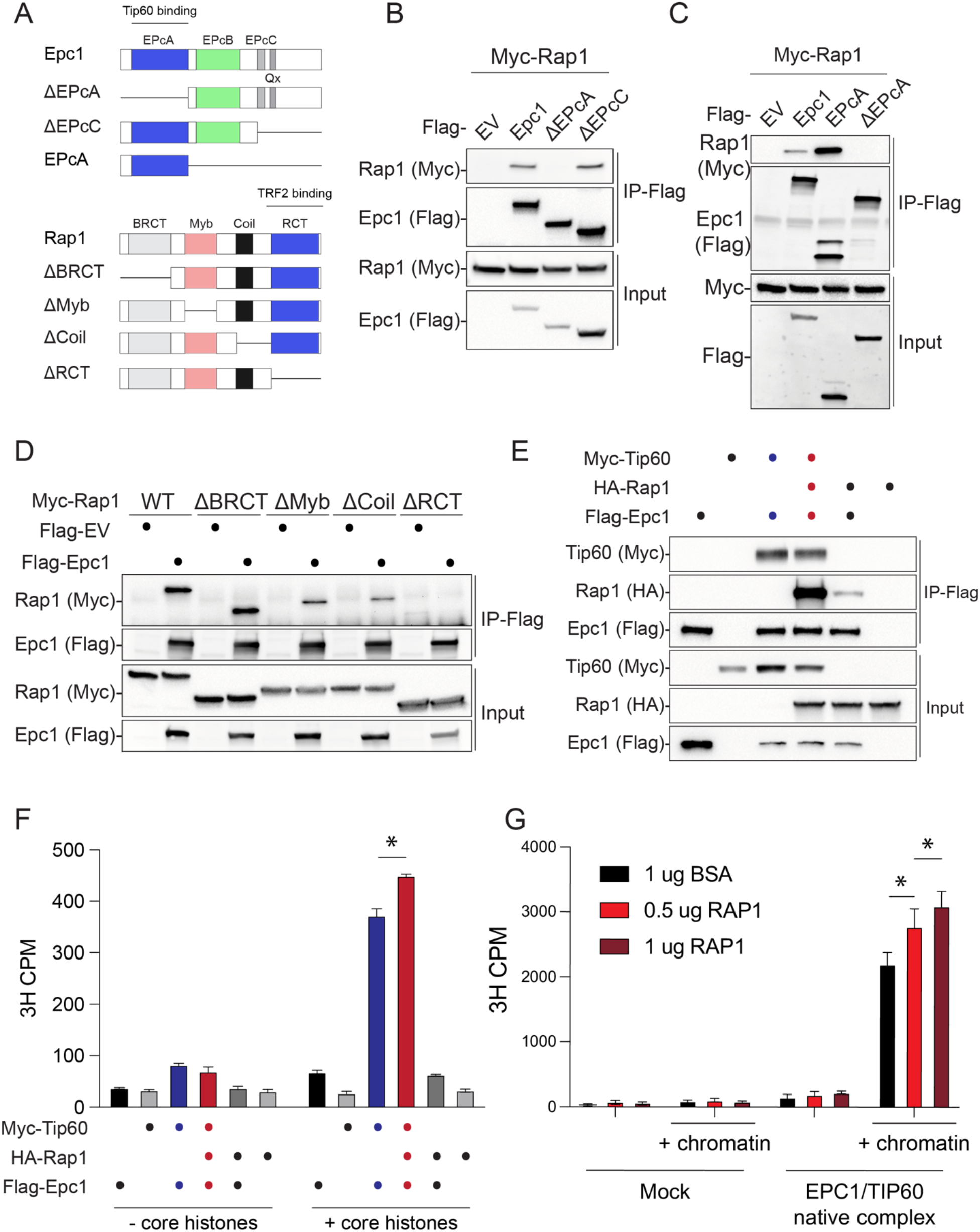
Rap1 forms a complex with and modulates the histone acetyltransferase (HAT) activity of Tip60/Epc1. A) Schematic illustration of Epc1 and Rap1 protein domains and deletion mutant constructs used for CoIP assays. B) CoIP of Flag-Epc1 deletion mutants and Myc-Rap1 in co-transfected HEK293T cells. Western blot analysis for Input and IP sample was performed with the indicated antibody. C) CoIP of Flag-Epc1 EPcA domain and Myc-Rap1. D) CoIP of Flag-Rap1 deletion mutants and Myc-Epc1. E) CoIP of Flag-Epc1, Myc-Tip60, and HA-Rap1. F) Histone acetyltransferase (HAT) activity assays performed using CoIPs from *E* on core histones. Data are the counts per minute (CPM) mean ± standard deviation of n = 2 technical replicates; two-way ANOVA, Tukey’s multiple comparison test; * p-value < 0.0001. G) HAT activity assay using affinity purified EPC1/TIP60 and Rap1 (increasing amounts) on short oligonucleosome chromatin. Bovine serum albumin (BSA) is a negative control. Data are CPM mean ± standard deviation of n = 3 technical replicates; two-way ANOVA, Tukey’s multiple comparison test; * p-value < 0.01.

We then explored whether RAP1, EPC1, and TIP60 form a trimeric complex. To that end, we co-expressed all three proteins in HEK293T cells and performed CoIP experiments. Using Rap1 as a bait, we found that Rap1-Epc1 and Rap1-Tip60 interactions were strongly enhanced in the presence of Tip60 and Epc1, respectively (Figure S4B). As expected, the effect was dependent on the EPcA domain of EPC1 (Figure S4C). In a parallel approach, we used EPC1 as bait and found that TIP60 greatly enhanced Rap1/Epc1 association (Figure 4E). Notably, histone acetyltransferase (HAT) assays revealed that Rap1 binds EPC1/TIP60 in an active conformation (Figure S4D) and increases Epc1/Tip60 HAT activity (Figure 4F-G). This was confirmed by CoIP of EPC1/TIP60/RAP1 from HEK293T cells (Figure 4F) and by *in vitro* addition of affinity purified human RAP1 protein to purified native TIP60/p400 complexes (Figure 4G). Altogether, we conclude that extratelomeric Rap1 associates with the TIP60/p400 complex and cooperatively binds to and modulates the HAT activity of Tip60/KAT5, thus illuminating the potential for Rap1 to act as a cofactor to regulate gene expression through histone modifications.

### Rap1 suppresses the 2-cell-like state in mouse embryonic stem cells

It has been established that the TIP60/p400 complex is essential for stem cell self-renewal and survival (Fazzio et al. 2008; Acharya et al. 2017), and members of the TIP60 complex (e.g. TIP60, DMAP1 and Ep400) suppress the emergence of “2-cell-like” (2C- like) cells in mouse embryonic stem cell cultures (mESC) (Rodriguez-Terrones et al. 2018). 2C-like cells were identified based on their resemblance to embryos at the two-cell stage of development (Macfarlan et al. 2012) and represent a unique window during preimplantation development when the zygotic genome becomes activated, marking a brief period of totipotency. During zygotic genome activation (ZGA), murine endogenous retrovirus with leucine tRNA primer (MERVL) elements are derepressed, along with the upregulation of several 2C genes such as *Zscan4* and *Dux.* Inner cell mass (ICM) derived mESC cultures contain a small fraction (<1%) of 2C-like cells exhibiting a 2C transcriptional state, providing a useful model to study totipotency (Macfarlan et al. 2012; Fu et al. 2019; Fu et al. 2020). Emergence of the 2C-like state is regulated by an atypical chromatin-assembly pathway involving FACT and CAF-1 histone chaperone complexes, the noncanonical PRC1 complex, and TIP60/p400 (Ishiuchi et al. 2015; Rodriguez-Terrones et al. 2018; Chen et al. 2020), although the exact molecular mechanisms are still to be determined.

The direct binding of RAP1 to members of the TIP60 complex (Figure 3D) prompted us to investigate whether RAP1 regulates the 2C-like state. To that end, we established independent mESC cultures from *Rap1^+/+^* and *Rap1^-/-^* mouse preimplantation blastocysts and performed RNA-seq. Transcriptomic analysis identified 202 upregulated genes (Figure 5A, Figure S5A; Supplemental Table S7), half of which (102; 50.5%) were previously categorized as genes expressed in 2C-like cells (Fu et al. 2020) (Figure 5A- B), including the *Zscan4* locus that we corroborated using RT-qPCR and IF (Figure S5B- C). We also observed that *Rap1^-/-^* mESCs displayed increased expression of the MERVL retrotransposon elements MT2_Mm, MERVL-int, and MERVL_2A-int, known to be de- repressed in 2C-stage embryos (Figure 5C and 5SD; Supplementary Table S8). Lastly, to address whether RAP1 suppresses the 2C-like state independent of telomere binding, we derived mESC cultures from *Rap1^I312R/I312R^* mouse blastocysts (Figure S5E) and found a similar expression of 2C-like genes in *Rap1^I312R/I312R^* mESCs relative to wild type (Figure S5F-G). In summary, our results uncover an unexpected function for a highly conserved telomere binding protein in the regulation of pluripotency.

**Figure 5.**
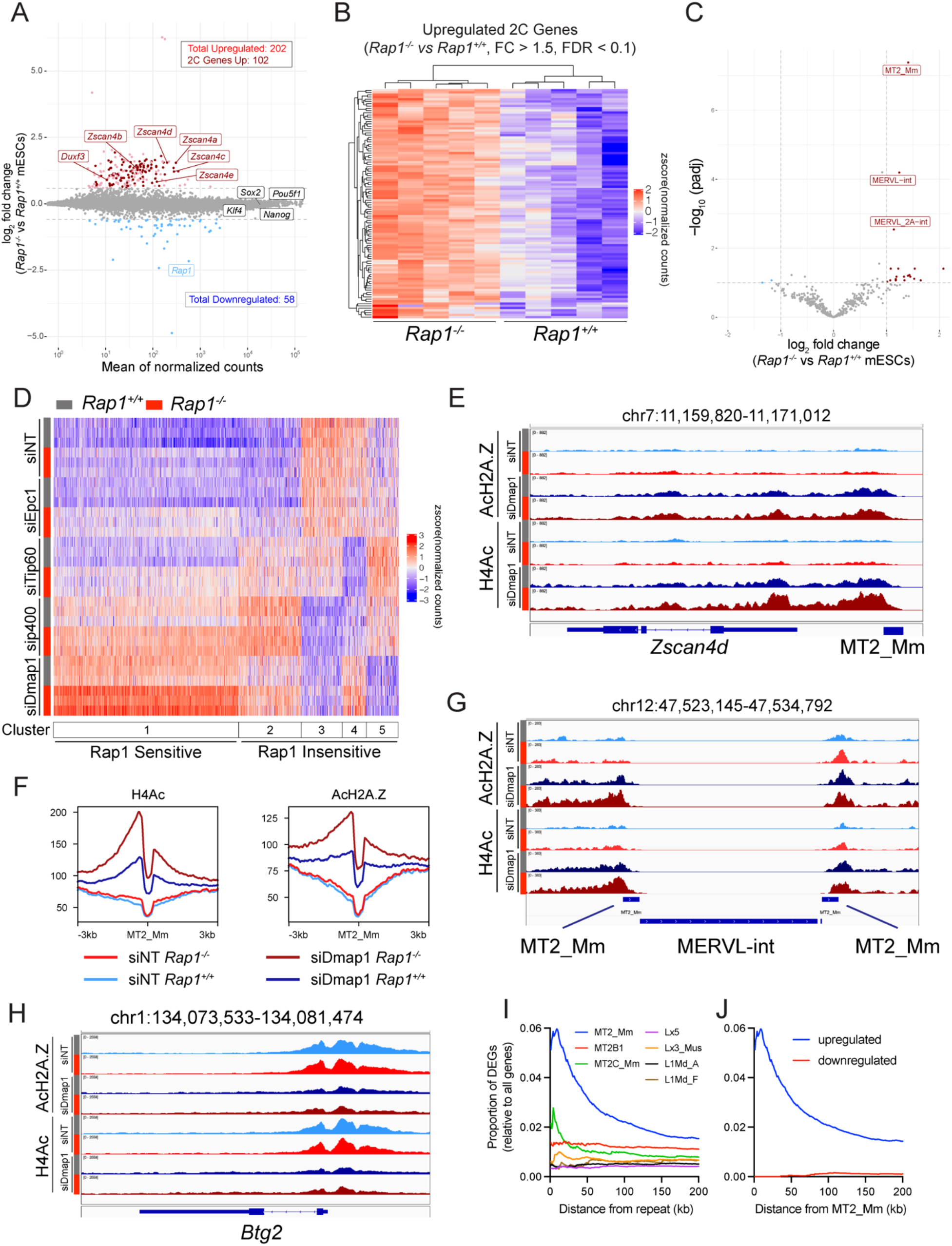
Rap1 suppresses the 2C-like state by enhancing Tip60/p400 repression of 2C genes. A) MA plot of log_2_ fold changes in gene expression in *Rap^-/-^ vs. Rap1^+/+^* mESCs. Upregulated (light red) and downregulated (light blue) genes are indicated (FC > 1.5; FDR < 0.1; n = 5 biological replicates). 2C-stage genes are highlighted in dark red. B) Hierarchically clustered heatmap representing RNA-seq data of 2C genes upregulated in *Rap^-/-^ vs. Rap1^+/+^* mESCs (n = 5 biological replicates). C) Volcano plot of differential repeat expression analysis in *Rap1^-/-^ vs. Rap1^+/+^* mESCs. Upregulated (red) and downregulated (blue) repeats are indicated (FC > 2; FDR < 0.1; n = 5 biological replicates). D) Kmeans (k = 5) clustered heatmap representing RNA-seq data of 2C genes in *Rap1^+/+^* (grey bar) and *Rap1^-/-^* (red bar) mESCs treated with siRNAs targeting Tip60/p400 subunits (Epc1, Tip60, p400, Dmap1) or nontargeting (siNT) control. 3 biological replicates were used per condition. E) ChIP-seq of H4Ac and AcH2A.Z profiles at the *Zscan4d* gene locus in *Rap1^+/+^* (grey bar) and *Rap1^-/-^* (red bar) mESCs transfected with siRNA targeting Dmap1 or nontargeting (siNT) control. Profiles plotted are representative of n = 3 biological replicates. F) Density plot centered at all MT2_Mm sites for H4Ac and AcH2A.Z ChIP-seq data. Density plots are representative of n = 3 biological replicates. G) ChIP-seq profiles of H4Ac and AcH2A.Z at a representative MERVL site in *Rap1^+/+^* (grey bar) and *Rap1^-/-^* (red bar) mESCs transfected with siRNA targeting Dmap1 and non-targeting control (siNT). H) ChIP-seq profiles of H4Ac and AcH2A.Z at a representative downregulated gene (*Btg2*) locus in *Rap1^+/+^* (grey bar) and *Rap1^-/-^* (red bar) mESCs transfected with siRNA targeting Dmap1 or nontargeting (siNT) control. I) Plot of the proportion of differentially expressed genes (DEGs) in *Rap1^+/+^ vs. Rap1^-/-^* mESCs (FC > 1.5; FDR < 0.1) at fixed distances from the indicated MERVL (MT2_Mm, MT2B1, MT2C_Mm) and LINE1 (Lx5, Lx3_Mus, L1Md_A, L1Md_F) repeat sequences. Proportions are calculated by dividing the number of DEGs by the total number of genes. J) Plot of the proportion of upregulated (blue) and downregulated (red) DEGs in *Rap1^+/+^* vs *Rap1^-/-^* mESCs (FC > 1.5; FDR < 0.1) at fixed distances from MT2_Mm repeat sequences.

### The interplay between RAP1 and the TIP60/p400 complex on mESC gene expression

Having established that extratelomeric RAP1 suppresses the 2C-like state in mESCs, we sought to better understand the interplay between Rap1 and Tip60/p400 during repression of 2C genes. We depleted Tip60, Epc1, Dmap1, and Ep400 individually by siRNA in *Rap1^+/+^* and *Rap1^-/-^* mESCs (Figure S6A), and RNA-seq analysis was performed. We first focused on the TIP60/p400 complex and interrogated the effect of depleting individual subunits on the expression of 2C-stage genes (Fu et al. 2020). Kmeans clustering highlighted 5 distinct 2C-state gene clusters (Figure 5D). Depletion of *Tip60/p400* subunits showed the expected reduction in transcription at three gene clusters (Figure 5D, clusters #3, #4, and #5). As for clusters #1 and #2, depletion of *Tip60*, *Ep400*, and *Dmap1* led to a paradoxical increase in 2C gene expression (Figure 5D, Clusters #1 and #2), suggestive of a repressive function for this complex in mESCs transcription (Fazzio et al. 2008; Rodriguez-Terrones et al. 2018). Of note, the various 2C gene clusters were not equally sensitive to inhibition of the various TIP60 members. For example, depletion of *Dmap1* and *Tip60* lead to opposite effects on gene expression within cluster #5. Furthermore, *Tip60* loss majorly impacted gene cluster #4, whereas *p400* and *Dmap1* depletion caused significant downregulation in gene expression for cluster #3. The latter observation can be explained by *Dmap1/p400*-mediated deposition of the histone variant H2A.Z as opposed to TIP60-driven histone acetylation. Overall, this sort of analysis revealed a high degree of complexity in how the TIP60/p400 complex regulates 2C-stage genes in mESCs, with certain subunits acting as negative repressors while others exerting a positive effect on gene transcription.

We next overlayed the impact of *Rap1* depletion on Tip60/p400 dysregulation of 2C gene expression and noted that half of all 2C genes (53.5%; 571 of 1068) were sensitive to *Rap1* loss (Figure 5D). Strikingly, RAP1-sensitive genes were further upregulated upon Tip60/p400 inhibition, including *Zscan4c* and MERVL (Figure S6B-C). Instead, gene clusters that were downregulated upon Tip60/p400 knockdown were largely insensitive to RAP1 (cluster #3, #4, and #5). Taken together, these results implicate RAP1 in the noncanonical and poorly understood function of TIP60/p400 during 2C genes supression.

### Increased histone acetylation at MERVL upon Rap1 loss

Examination of the *Zscan4* locus and the MERVL LTR MT2_Mm by ChIP-seq revealed increased H4Ac and AcH2A.Z abundance in *Rap1^-/-^* cells, that was further enhanced upon depletion of Dmap1 (Figure 5E-G). As a control, we show that Rap1 insensitive genes did not exhibit differences in H4Ac or AcH2A.Z levels between *Rap1^+/+^* and *Rap1^-/-^* mESCs. Furthermore, genes that were downregulated upon Dmap1 depletion showed the expected decrease in H4Ac and AcH2A.Z at promoter regions (Figure 5H). Strikingly, a significant fraction of genes proximal to MERVL MT2_Mm elements were upregulated in *Rap1^-/^*^-^ mESCs (Figure 5I-J), suggesting that a significant proportion of genes differentially expressed upon Rap1 loss were at fixed distances from this class of repeat elements. We found no proximal distance association between upregulated genes and unrelated retroviral elements such as LINE1 (Lx5, Lx3_Mus, L1Md_A, L1Md_F) (Figure 5I). This implies that upregulation of a subset of 2C genes in the absence of Rap1 can be ascribed to the induction of genes proximal to MERVL. It has been reported that in 2C-like cells MERVL elements can serve as enhancers and create sites of cryptic transcription initiation, influencing proximal genes (Macfarlan et al. 2012). Accordingly, the paradoxical de-repression of 2C genes upon Tip60/p400 and Rap1 depletion could be the result of cis-effects of increased MERVL repeat elements expression.

## Discussion

Our study sheds light on the mechanism by which Rap1 regulates gene expression in mammals by establishing that RAP1 maintain native transcription independent of its association with telomeres. We find that extratelomeric RAP1 responsible for its transcriptional function does not bind to discrete genomic DNA loci. Instead, this conserved telomere binding protein modulates the HAT activity of the Tip60/p400 complex. Furthermore, we uncover an unanticipated role for mammalian RAP1 in regulating the repressive activity of the TIP60/p400 complex on 2C-stage genes and endogenous retroviruses in mESCs.

### Transcriptional regulation: an ancestral function for the most conserved telomere binding protein

RAP1 is the most highly conserved telomere binding protein and in unicellular eukaryotes serves an essential function in telomere maintenance by regulating telomere length, preventing telomere fusions, and suppressing telomere-telomere recombination events (Kanoh and Ishikawa 2001; Pardo and Marcand 2005; Nanavaty et al. 2017). When compared to its chimpanzee counterpart, the gene encoding human RAP1 shows little divergence (0.25 aa changes/100 aa) relative to other shelterin subunits (0.65 aa changes/100 aa for TRF2, TRF1, and TIN2, respectively) (Kabir et al. 2014). Yet, RAP1 is largely dispensable for telomere function and cell viability in mammals (Sfeir et al. 2010; Kabir et al. 2014). This paradox raises a question related to the selective pressure for RAP1 conservation in higher eykaryotes. We speculate that the function of RAP1 in gene regulation might underly such conservation. In budding yeast, Rap1 binds to DNA sequences at gene promoters and is necessary to maintain high expression of genes encoding ribosomal proteins and glycolytic enzymes, without which cell growth and proliferation are greatly stunted (Bram and Kornberg 1985; Huet et al. 1985; Shore and Nasmyth 1987; Vignais et al. 1987; Chambers et al. 1989). Additionally, in both yeast and *T. brucei*, Rap1 maintains repression of telomere/subtelomere regions, preventing aberrant expression of genes proximal to chromosome ends, such as the variant surface glycoprotein (VSG) genes found in *T. brucei* subtelomeres (Yang et al. 2009). Rap1 transcriptional activity in budding yeast is also linked to telomeres. Specifically, its binding to telomeres represses the transcription of telomeric repeat-containing RNA (TERRA) – a long non-coding RNA in subtelomere regions – by recruiting of Rif1/2 and Sir2/3/4 complexes (Iglesias et al. 2011). Furthermore, as telomeres shorten in budding yeast, the abundance of extratelomeric RAP1 increases, facilitating its localization to additional target genes (called NRTS, for new Rap1 targets at senescence) to regulate transcription (Platt et al. 2013; Song et al. 2020). In contrast, our study shows that mammalian RAP1 transcriptional activity is uncoupled from telomere binding, with independent pools of RAP1 associating with either TRF2 or TIP60. We show that mutant RAP1 (Rap1^I312R^) that cannot bind TRF2 or localize to telomeres fully complements gene expression (Figure 1G). Furthermore, whereas yeast Rap1 binds directly to DNA sequences at gene promoters to regulate transcription, we found that the extratelomeric pool of mammalian RAP1 does not bind genomic loci (Figure 2D-E) and mammalian Rap1 lacks DNA binding activity (Figure 2G, S2G-H). Thus, although Rap1 function in transcription regulation is conserved in higher eukaryotes, the mechanism by which Rap1 regulates transcription has diverged.

### RAP1 as a regulator of the TIP60 complex

We establish that extratelomeric RAP1 associated with TIP60/p400, a histone acetyltransferase complex with a fundamental role in the regulation of gene transcription, as demonstrated by proximity labeling as well as co-immunoprecipitation with the Tip60, Epc1/Epc2, and Dmap1 subunits of the complex (Figure 3E). RAP1 binds to the Epc1 N-terminal EPcA domain through its C-terminal RCT domain (Figure 4). TRF2 also binds Rap1-RCT and appears to compete with EPC1 (Figure S4A), suggesting that RAP1 binding to TIP60/p400 is mutually exclusive with TRF2. This is consistent with the observation that Rap1 maintains native transcription independent of TRF2 and provides evidence that RAP1 associates with multiple regulatory protein complexes through its RCT domain. This is similar to budding yeast, where Rap1 utilizes its RCT domain to recruit a diverse set of regulatory factors depending on the genomic and functional context, for example, Rif1/Rif2 at telomeres to regulate telomere length (Hardy et al. 1992b; Wotton and Shore 1997) and Sir3/Sir4 at *HML/HMR* to maintain transcriptional silencing (Moretti et al. 1994; Moretti and Shore 2001). Association of RAP1 with the TIP60/p400 complex is conserved between humans and mice, as demonstrated by a recent publication that used a similar proximity-dependent biotinylation approach in human cells (Go et al. 2021). In addition, our study shows RAP1 modulates TIP60/p400 HAT activity (Figure 4F-G), establishing RAP1 as a transcriptional cofactor within a chromatin modifying complex. Future structural and biochemical studies are necessary to understand how Rap1 regulates the activity of the Tip60/p400 complex.

Importantly, we found that loss of RAP1 synergizes with Tip60/p400 depletion to increase H4 and H2A.Z acetylation at promoters of upregulated 2C genes (Figure 5E-F). The noncanonical activity of TIP60/p400 on mESC gene expression has been previously noted (Fazzio et al. 2008; Chen et al. 2013) and contrasts the case in somatic cells, where TIP60/p400 is primarily a transcriptional activator that leads to histone acetylation. So far, the underlying molecular mechanism of TIP60 mediated repression of gene expression remains completely unknown, and our data implicated RAP1 in the non canonical function of TIP60. Furthermore, our results suggest that the majority of RAP1 dysregulated genes are proximal to MERVL repeat elements which also exhibit elevated H4/H2A.Z acetylation (Figure 5G-I).This proximity-dependent effect on gene expression has also been observed upon depletion of the FACT and CAF-1 histone chaperone complexes, contexts in which cryptic transcription generates MERVL-fused chimeric transcripts (Ishiuchi et al. 2015; Chen et al. 2020). Taken together, we propose a model whereby the activation of endogenous retroviral elements MERVL induces the transcription of proximal genes in context of Rap1 deficiency. However, we cannot exclude that the effect of Rap1 on 2C genes is downstream of Zscan4 activation (Zhang et al. 2019) or perhaps due to differential acetylation of non-histone proteins which is another known mechanism by which Tip60/p400 modulates gene expression (Sapountzi and Cote 2011).

### Telomere binding proteins and totipotency

An unlikely link between telomeres, telomere binding proteins, and pluripotency emerged in recent literature. Loss of TRF1 in stem cells leads to increased TERRA levels that in turn modulated the recruitment of the PRC2 repressive complex to alter pluripotency genes expression (Marion et al. 2019). More recently, depletion of TRF2 in mESCs was shown to induce the expression of 2C-stage genes, including *Zscan4* which contributed to telomere protection in the absence of TRF2 (Markiewicz-Potoczny et al. 2021). The latter study highlights an unexpected link between totipotency and telomere protection mechanisms. Our study unravels yet another connection between telomere biology and totipotency, as we find that RAP1 suppresses a 2C-like state in mESCs by enhancing the repressive activity of TIP60/p400 on MERVL and other 2C genes (Figure 5A-D). In effect, our study highlights how the telomere binding protein Rap1 interfaces with chromatin modifying complexes in the control of totipotency. It is tempting to speculate that the regulation of transcription at retroviral repeat elements constitute an ancestral function for RAP1, and possibly other shelterin subunits prior to being co-opted as a telomere binding proteins. Finally, further studies are necessary to explore the impact of this telomere binding protein on other classes of endogenous retrovirus and in different biological settings.

## Acknowledgements

We thank members of the Sfeir lab for feedback and comments on the manuscript. pRK5- HA-Ubiquitin-K48 was a gift from Ted Dawson (Addgene, #17605). We are thankful to Karim-Jean Armache for guidance on RAP1 gel shift assays. Catherine Lachance is acknowledged for purified NuA4/TIP60 and chromatin fractions. We acknowledge Beatrix M. Ueberheide and the Proteomics lab as well as Sang Y. Kim and the Rodent Genetic Engineering lab at NYU School of Medicine. We thank the Genome Technology Center at NYU School of Medicine and the Integrated Genomics Operation at Sloan Kettering Institute, MSKCC. This work was supported by a grant from the NIH (R01 DK102562) to A.S. and CIHR (FDN-143314) to J.C. R.M.B. is supported by a training grant from NIH (1F30 DK118901). C.H. lab is funded by internal institutional support of the Institute of Biophysics of the Czech Academy of Sciences (68081707), the Czech Science Foundation, project 19-18226S, and Ministry of Education, Youth and Sports of the Czech Republic. project LTAUSA19024

## Authors Contribution

A.S., R.M.B. conceived the experimental design. R.M.B. performed experiments with the help of P.S., O.S., and W.K. M.A. in C.H. lab performed EMSA and RAP1 purification.

A.M. in J.C. lab performed HAT assays. P.D. prepared nucleosomes for EMSA. A.S. and

R.M.B. wrote the manuscript. All authors discussed the results and commented on the manuscript.

## Author information

Agnel Sfeir is a co-founder, consultant, and shareholder in Repare Therapeutics.

Correspondence and requests for materials should be addressed to A.S.

## Supplemental Figures

**Figure S1.**
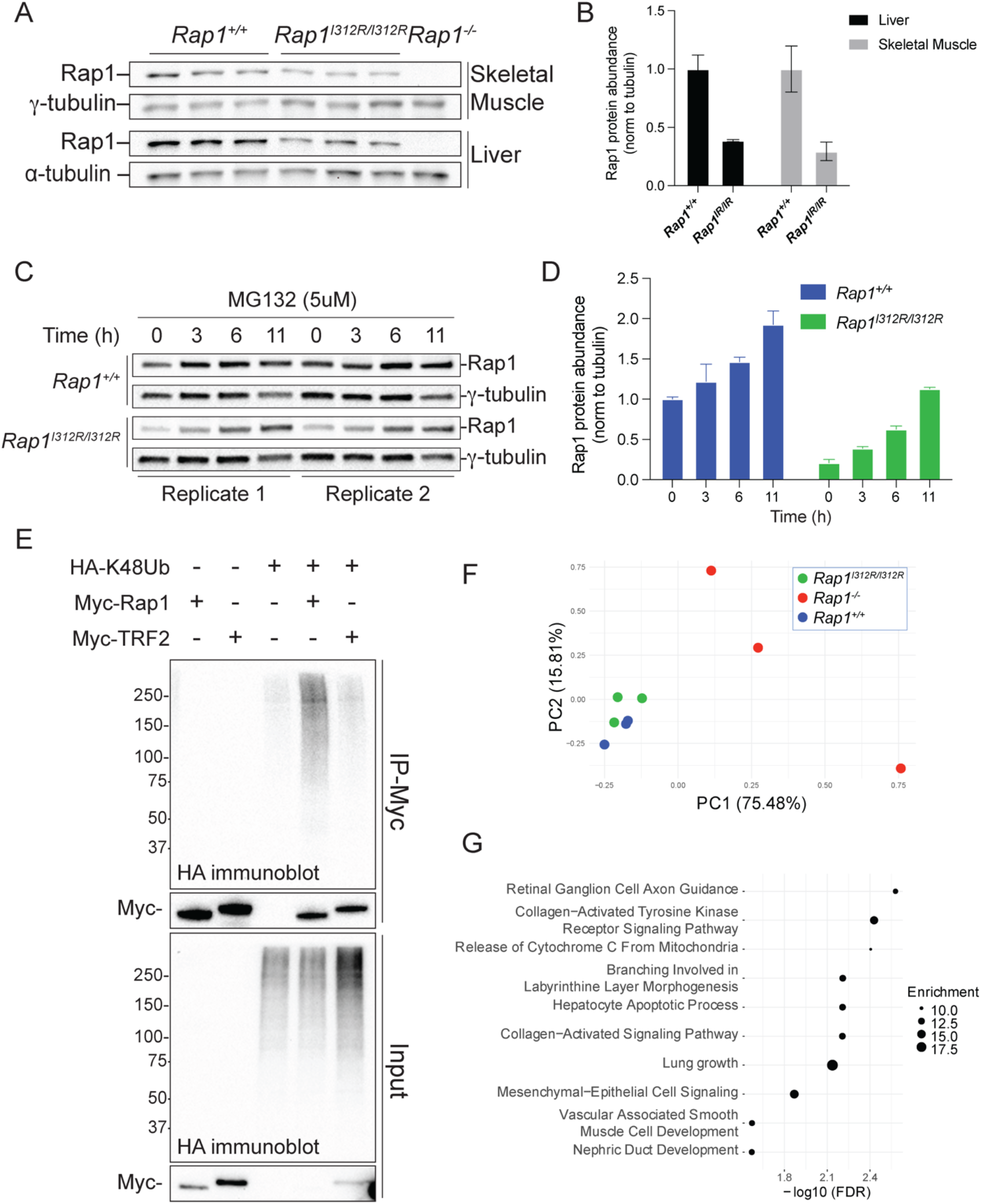
related to Figure 1: Analysis of extratelomeric Rap1 protein level regulation and impacts on gene regulation. A) Immunoblot for Rap1 from *Rap1^-/-^* (n = 1), *Rap1^+/+^* (n = 3), and *Rap1^I312R/I312R^* (n = 3) mouse skeletal (gastrocnemius) muscle and liver lysates. *γ*- or *α*-tubulin are used as loading controls. B) Quantification of Rap1 levels in (A) Rap1 relative abundance was determined by normalizing to *γ*- or *α*-tubulin, as indicated. Data is the mean ± standard deviation of n = 3 biological replicates. C) Immunoblot for Rap1 from *Rap1^+/+^* and *Rap1^I312R/I312R^* MEFs (n = 2 biological replicates) following treatment with proteasome inhibitor MG132 (5uM) for 11 hours. *γ*-tubulin is used as loading control. D) Quantification of Rap1 levels in C. Rap1 relative abundance was determined by normalizing to *γ*-tubulin. Data is the mean ± standard deviation of n = 2 biological replicates. E) CoIP of Myc-tagged Rap1, TRF2, and HA-K48-linked ubiquitin (K48Ub) in co-transfected HEK293T cells. F) Principal component analysis (PCA) of RNA-seq data from *Rap1^+/+^, Rap1^I312R/I312R^*, and *Rap1^-/-^* MEFs. Percent along axes indicate the proportion of variance due to the indicated principal component. G) Graphical representation of top 10 biological processes overrepresented in differentially expressed genes (DEGs) between *Rap1^+/+^* and *Rap1^-/-^* MEFs (FC > 1.5; FDR < 0.1) using PANTHER Classification System statistical overrepresentation test.

**Figure S2.**
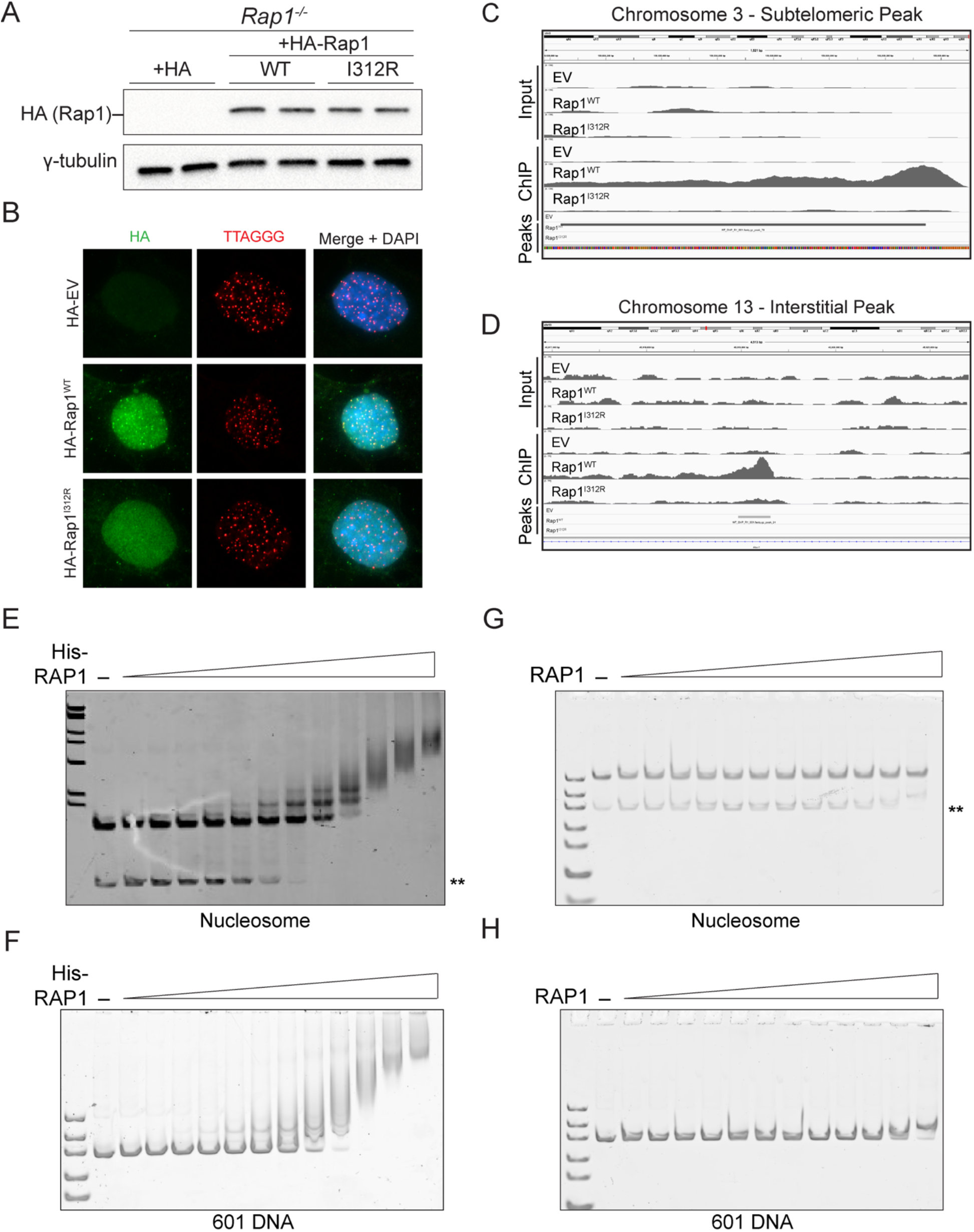
related to Figure 2: Analysis of Rap1 chromatin and DNA binding activity using ChIP-seq and EMSA. A) Immunoblot for Rap1 (anti-HA antibody) in *Rap1^-/-^* MEFs expressing HA-Rap1-WT, HA-Rap1-I312R, or empty vector (EV) control (n = 2 biological replicates). *γ*-tubulin is used as loading control. B) Representative IF-FISH image in *Rap1^-/-^* MEFs expressing HA-Rap1-WT, HA-Rap1-I312R, and empty vector (EV) control and stained for Rap1 (anti-HA antibody green) and telomeres (TTAGGG PNA probe in red). DAPI (blue) is used as counterstain. C) HA-ChIP-Seq profile at representative HA-Rap1- WT subtelomeric peak on chromosome 3. Profiles plotted are representative of n = 4 biological replicates. D) HA-ChIP-Seq profile at representative HA-Rap1-WT interstitial peak on chromosome 13. E) Electrophoretic mobility shift assay (EMSA) using His-tagged Rap1 and affinity purified nucleosome substrate. **free DNA found in nucleosome preparations. F) EMSA using non-His-tagged Rap1 and purified nucleosome. G) EMSA using His-tagged Rap1 and double-stranded 601 DNA substrate. H) EMSA using non-His-tagged Rap1 and double-stranded 601 DNA substrate.

**Figure S3.**
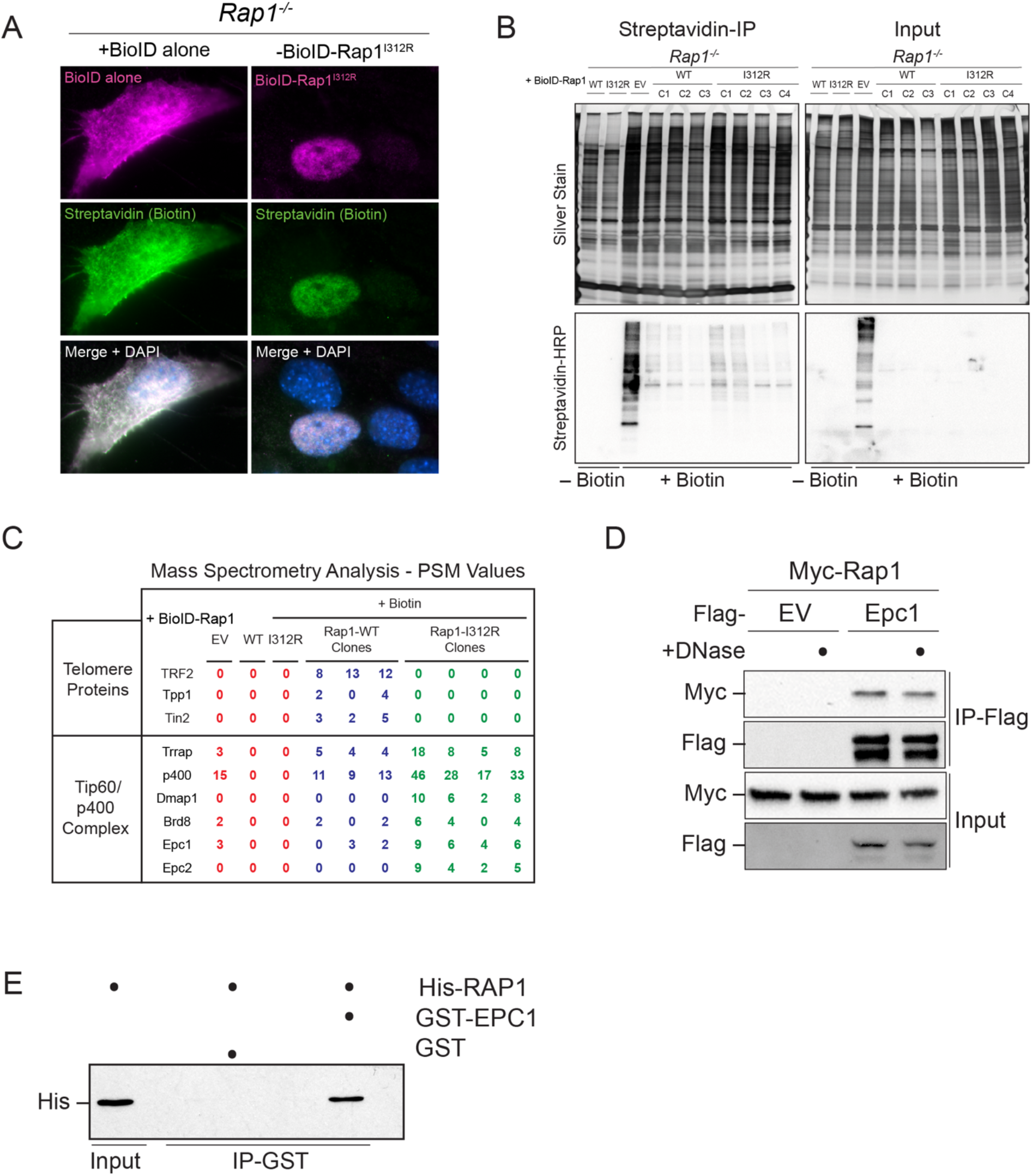
related to Figure 3: Analysis of Rap1 protein interactors by proximity-dependent biotinylation. A) Representative images of streptavidin-IF in *Rap1^-/-^* MEFs expressing BioID-Rap1-I312R and empty vector (EV) control for BioID (anti-FLAG; magenta) and biotin (streptavidin antibody in green). DAPI (blue) is used as counterstain. B) Silver stain (top) and Western blot for streptavidin-HRP (bottom) following streptavidin pull-down in *Rap1^-/-^* MEFs expressing BioID-Rap1-WT (n = 3 biological replicates; clones C1, C2, C3), BioID-Rap1-I312R (n = 4 biological replicates; clones C1, C2, C3, C4), and BioID alone (EV). Where indicated, cells were treated with biotin (50 μM) for 24 hours prior to harvest. C) Summary of peptide-spectrum match (PSM) values of telomere proteins (top) and Tip60/p400 complex members (bottom) from BioID mass spectrometry analysis. D) CoIP of Flag-Epc1 and Myc-Rap1 following co-transfection of HEK293T cells. Where indicated, lysates were treated with DNase (10 μg/mL) for 1 hr at room temperature prior to incubation with antibody for IP. E) Pull-down of GST-EPC1 and His- RAP1 using glutathione-coupled beads. GST tag alone is used as negative control. Blot is representative of n = 2 technical replicates.

**Figure S4.**
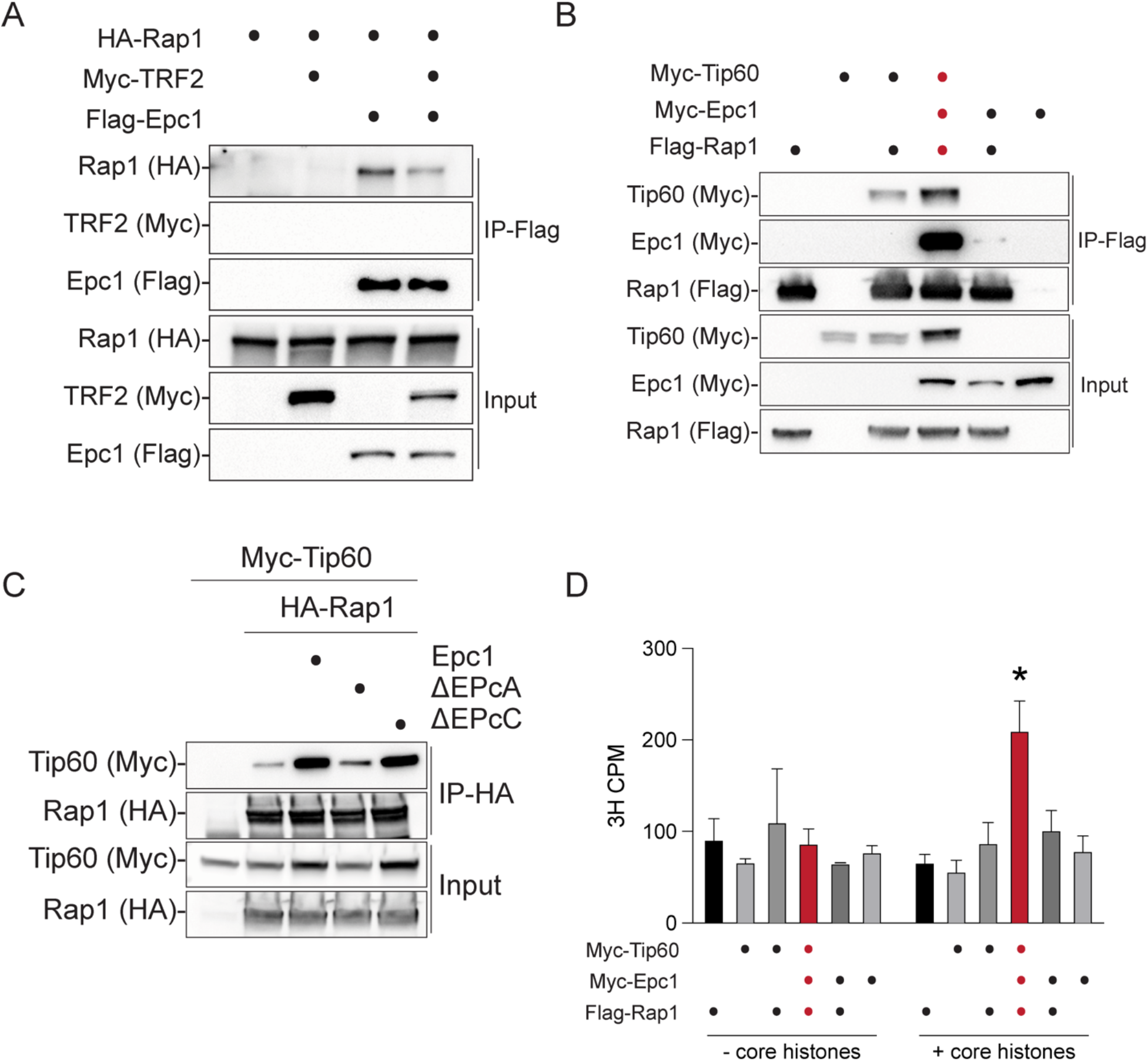
related to Figure 4: Biochemical analysis of Rap1 binding to Epc1 and Tip60. A) CoIP of Flag-Epc1 and HA-Rap1 +/- Myc-TRF2 from HEK293T cells cotransfected with the indicated plasmid. B) CoIP of Flag-Rap1, Myc-Epc1, and Myc- Tip60. C) CoIP of HA-Rap1, Myc-Tip60, and Epc1 deletion mutants. D) Histone acetyltransferase (HAT) activity assays using CoIPs from *B* and core histones. Data presented as the counts per minute (CPM) mean ± standard deviation of n = 2 technical replicates; two-way ANOVA, Tukey’s multiple comparison test; * p-value < 0.01.

**Figure S5.**
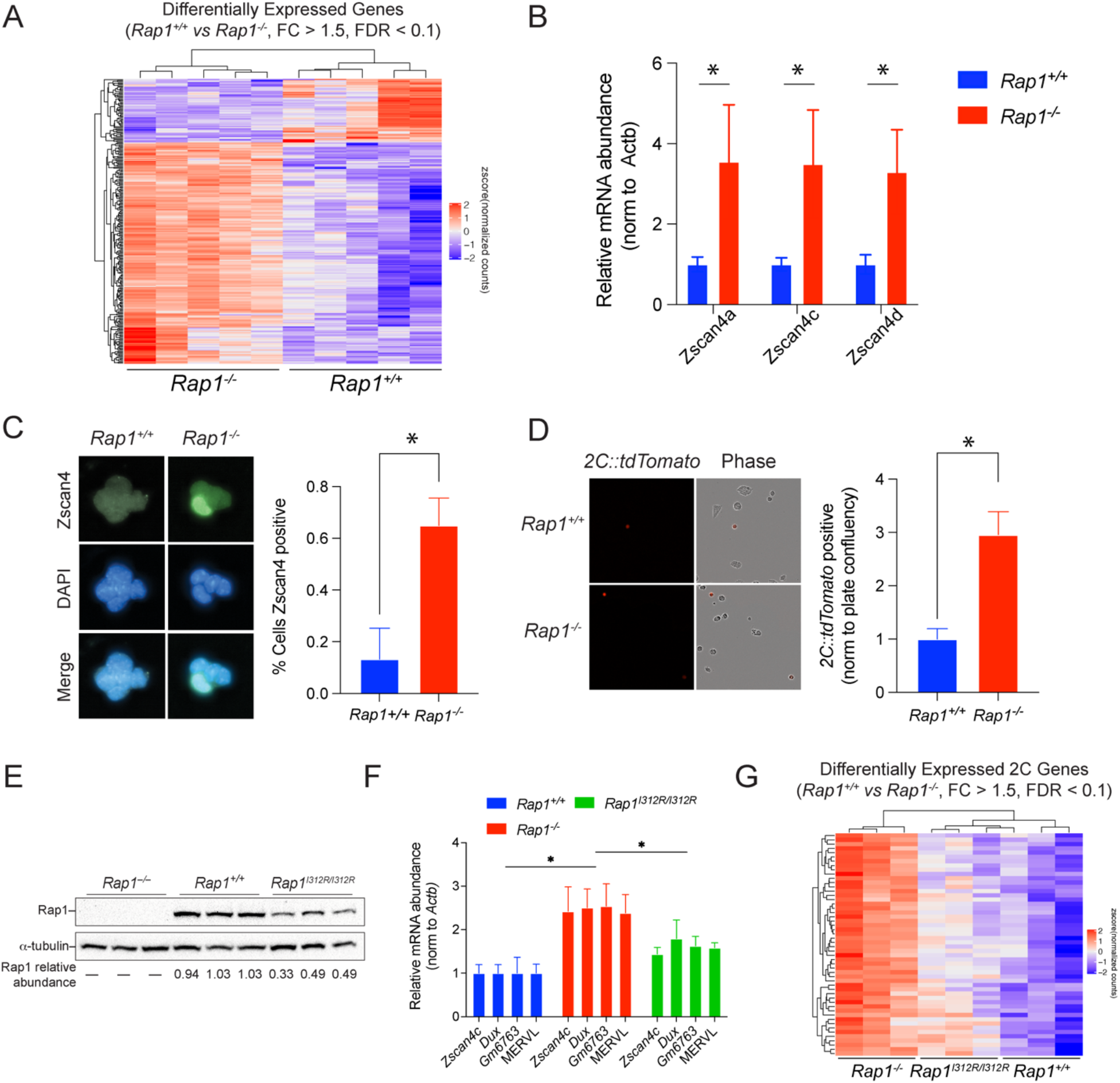
related to Figure 5: Analysis of transcriptomes and 2C-like state in *Rap1^+/+^*, *Rap1^-/-^*, and *Rap1^I312R/I312R^* mESCs. A) Hierarchically clustered heatmap representing RNA-seq data for differentially expressed genes (DEGs) between *Rap1^+/+^* and *Rap1^-/-^* mESCs (FC > 1.5; FDR < 0.1; n = 5 biological replicates per genotype). B) qRT-PCR for *Zscan4* locus (*Zscan4a, Zscan4c, Zscan4d*) in *Rap1^+/+^* and *Rap1^-/-^* mESCs. Relative mRNA abundance is determined by normalizing to *ACTB* using the delta-delta Ct method. Data are mean ± standard deviation of n = 3 biological replicates; student’s T test; * p-value < 0.05. C) (Left) Representative images of IF for Zscan4 (green) in *Rap1^+/+^* and *Rap1^-/-^* mESCs. DAPI (blue) is used as counterstain. (Right) Percent of Zscan4 positive cells in *Rap1^+/+^* and *Rap1^-/-^* mESCs. Data are mean ± standard deviation of n = 3 biological replicates; student’s T test; * p-value < 0.01. D) (Left) Representative images of live-cell imaging for *2C::tdTomato* reporter (red) in *Rap1^+/+^* and *Rap1^-/-^* mESCs. Phase contrast imaging is used to determine cell confluency. (Right) Relative number of *2C::tdTomato* positive cells in *Rap1^+/+^* and *Rap1^-/-^* mESCs normalized to confluency. Data are mean ± standard deviation of n = 3 biological replicates; student’s T test; * p-value < 0.01. E) Immunoblot for Rap1 from *Rap1^-/-^*, *Rap1^+/+^*, and *Rap1^I312R/I312R^* mESC whole cell lysates (n = 3 biological replicates). Rap1 relative abundance was determined by normalizing to *α*-tubulin. F) qRT-PCR for 2C genes (*Zscan4c, Dux, Gm6763*) and MERVL in *Rap1^+/+^*, *Rap1^-/-^*, and *Rap1^I312R/I312R^* mESCs. Relative mRNA abundance is determined by normalizing to *ACTB* using the delta-delta Ct method. Data are mean ± standard deviation of n = 3 biological replicates; 2-way ANOVA, Dunnett’s multiple comparison test; * p-value < 0.05. G) Hierarchically clustered heatmap representing RNA-seq data for differentially expressed 2C genes between *Rap1^+/+^* and *Rap1^-/-^* mESCs (FC > 1.5; FDR < 0.1; n = 3 biological replicates).

**Figure S6.**
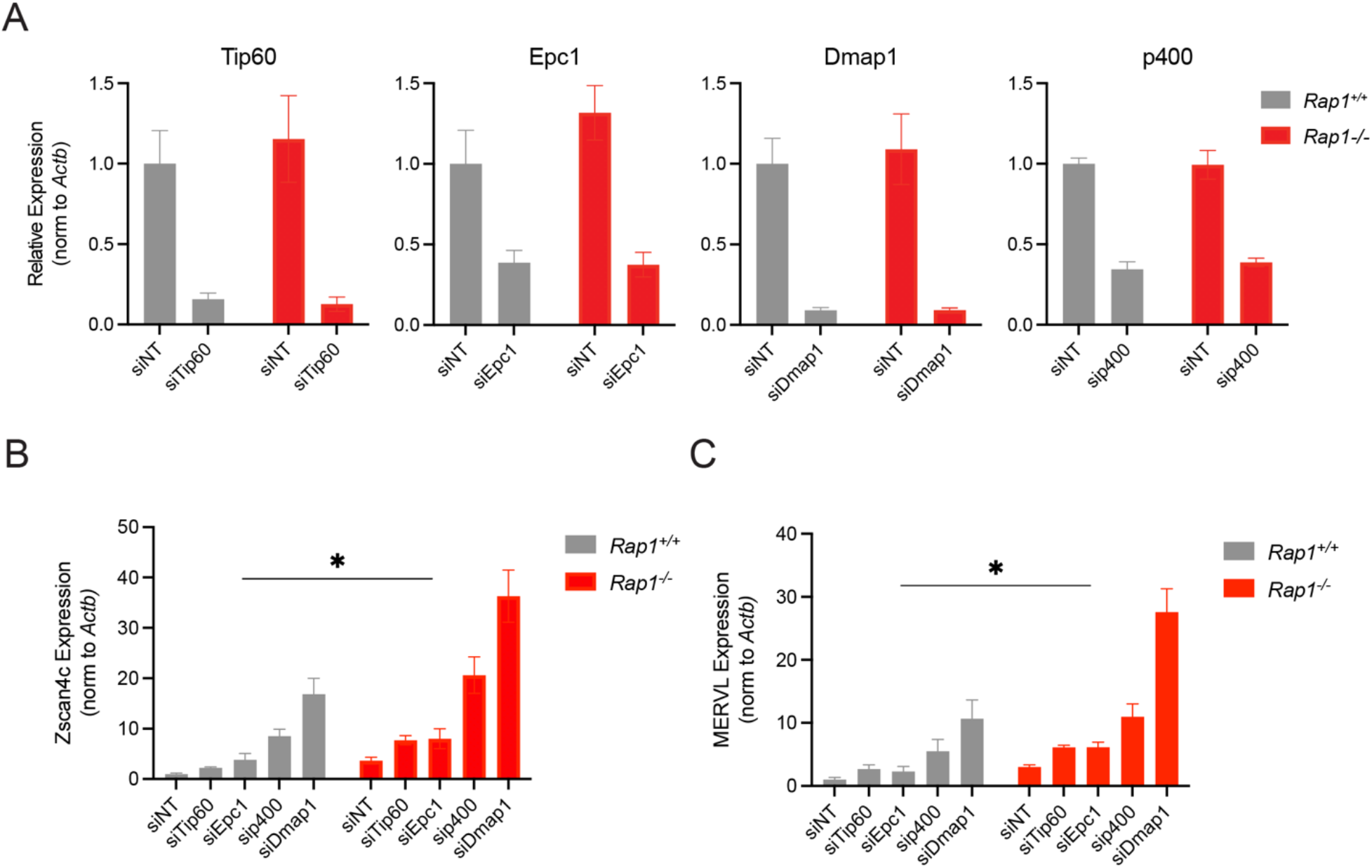
related to Figure 5: Investigating the interplay between Rap1 and Tip60/p400 on 2C gene transcription. A) qRT-PCR for the indicated gene in *Rap1^+/+^* and *Rap1^-/-^* mESCs treated with the indicated siRNA or nontargeting (siNT) control. Relative mRNA abundance is determined by normalizing to *ACTB* using the delta-delta Ct method. Data are mean ± standard deviation of n = 3 biological replicates. B) qRT-PCR for *Zscan4c* in *Rap1^+/+^* and *Rap1^-/-^* mESCs treated with the indicated siRNA or nontargeting (siNT) control. Relative mRNA abundance is determined by normalizing to *ACTB* using the delta-delta Ct method. Data are mean ± standard deviation of n = 3 biological replicates; student’s T test; * p-value < 0.05. C) qRT-PCR for MERVL in *Rap1^+/+^* and *Rap1^-/-^* mESCs treated with the indicated siRNA or nontargeting (siNT) control. Relative mRNA abundance is determined by normalizing to *ACTB* using the delta-delta Ct method. Data are mean ± standard deviation of n = 3 biological replicates; student’s T test; * p-value < 0.05.

## Methods

### Animal Studies

*Rap1* knockout mice were generated by crossing the *Rap1* floxed allele to *Rosa*-Cre mice, as previously described (Sfeir et al. 2010). *Rap1^I312R^* knock-in mice were generated by CRISPR/Cas9 editing via injection of fertilized oocytes with Cas9 mRNA, sgRNA targeting the *Rap1* locus (5’-TAGCTGCCGGATCACCTTAA-3’), and single-stranded DNA donor (ssODN) carrying the mutated allele (Supplementary Table S6). Two mice heterozygous for the mutation were obtained using this method and were crossed to produce homozygous mice. Genotyping was performed using the following primers: Fwd: 5’- CTCTCACACACACACCATGCATTC-3’; Rev: 5’-GACTCTAAGAAGGAGGACGTGG-3’.

AciI restriction digest produces a 734bp band for the Rap1-WT allele and two bands at 636bp and 98bp for Rap1-I312R.

For mouse embryonic fibroblast (MEF) isolation, pregnant mice were sacrificed at E13.5 and embryos were harvested and dissected to remove limbs, liver and neural tissue. The remaining tissue was diced, treated with trypsin, and then cultured in serum-containing media. Mouse embryonic stem cell (mESC) cultures were derived from the inner cell mass of preimplantation embryos.

### Plasmids

Mouse Rap1 variants (wild type, Rap1-I312R) were cloned into the retroviral constructs pLPC-N-Flag-2xHA-MCS-Puro for ChIP-seq and pLPC-N-Flag-BirA*-13xGGGGS-Myc- MCS-Puro for proximity-based biotin labeling. For CoIP, Rap1 variants (wild type, Rap1- I312R, Rap1-*Δ*BRCT, Rap1-*Δ*Myb, Rap1-*Δ*Coil Rap1-*Δ*RCT) were subcloned into the retroviral constructs pLPC-N-Myc-MCS, pLPC-N-Flag-MCS, and pcDNA-N-3xHA-MCS. Mouse Tip60, Epc1, Epc2, Dmap1, Ruvbl1, and Actl6a cloned into pCMV6-MCS-Myc-Flag were purchased from Origene and used for CoIPs in Figure 3E-G. Mouse Tip60 and Epc1 variants (wild type, Epc1-*Δ*EPcA, Epc1-*Δ*EPcC, Epc1-EPcA) were subcloned by restriction enzyme digest of PCR products from pCMV6-MCS-Myc-Flag into pLPC-N-Flag-MCS and pLPC-N-Myc-MCS for CoIPs in Figure 4B-E. Human RAP1 was expressed from pTriEx4 for protein affinity purification used in Figure 4G. For K48-linked ubiquitin CoIP, pRK5-HA-Ubiquitin-K48 was a gift from Ted Dawson (Addgene, #17605)

### Cell Culture

Primary MEFs were cultured in 15**%** fetal bovine serum (FBS), 1x MEM Non-Essential Amino Acids (Gibco 11140050), 100 U/mL Pen/Strep (Gibco 15140122), 2mM L-glutamine (Gibco 25030149), and 50 µM 2-mercaptoethanol in DMEM. SV40LT- immortalized MEFs were immortalized by retroviral infection with pBabeSV40LT (a gift from Greg Hannon) and cultured in 10**%** bovine calf serum (BCS), 1x MEM Non-Essential Amino Acids (Gibco 11140050), 100 U/mL Pen/Strep (Gibco 15140122), and 2mM L-glutamine (Gibco 25030149) in DMEM. mESCs were grown on feeders or on gelatinized plates in 2i/LIF media composed of 15% FBS, 1x NEAA, 100U/mL Pen/Strep, 2mM L-glutamine, 50 µM 2-mercaptoethanol, 500 U/mL LIF Protein (Millipore Sigma ESG1106), 3 µM CHIR99021/GSK-3 Inhibitor (Millipore Sigma 361571), and 1 µM PD0325901/MEK1/2 Inhibitor (Millipore Sigma 444968) in DMEM. Cells were tested for mycoplasma using the LookOut mycoplasma PCR detection kit (Sigma MP0035), following the manufacturer’s instructions and using the JumpStart Taq DNA polymerase

(Sigma D9307). For proteasome inhibitor studies, cells were treated with 5uM MG132 (Invivogen TLRL-MG132) or with dimethyl sulfoxide (DMSO) as control.

### Western Blot Analysis

Cells or tissues were either lysed directly in Laemmli buffer (Biorad 1610747) or in RIPA buffer (25mM Tris pH 7.6, 150mM NaCl, 1% NP-40, 1% sodium deoxycholate, 0.1% SDS), sonicated to shear chromatin and spun at high-speed to pellet cellular debris. If lysed in RIPA, protein concentration was determined by BCA assay (Thermo Scientific 23225). Equal amounts of protein were separated by SDS-PAGE, transferred to nitrocellulose and probed with antibody. After incubation with secondary antibody (Millipore Sigma GENA931, mouse; GENA934, rabbit), immunoblots were developed with enhanced chemiluminescence (Biorad 1705060). Protein abundance was quantified by measuring the integrated intensity of bands using Fiji ImageJ and normalizing to loading controls. The following primary antibodies were used: mouse Rap1 (1252, rabbit polyclonal), TRF2 (Novus Biologicals NB110-57130), *γ*-tubulin (Millipore Sigma T6557, mouse monoclonal), *α*-tubulin (Abcam ab7291, mouse monoclonal), histone H3 (Abcam ab1791, rabbit polyclonal), Flag (Cell Signaling #14793, rabbit monoclonal), c-Myc (Santa Cruz Biotech. sc-789, rabbit polyclonal), HA (Abcam ab9110, rabbit polyclonal), and streptavidin-HRP (Invitrogen 1953050).

### IF-FISH

MEFs and mESCs grown on coverslips were fixed with 4% paraformaldehyde for 10 min, washed with PBS, and permeabilized with 0.5% Triton X-100 buffer (0.5% Triton X-100, 20mM HEPES pH 7.9, 50mM NaCl, 3mM MgCl_2_, 300mM sucrose. Cells were then incubated in blocking solution (1mg/mL BSA, 3% goat serum, 0.1% Triton X-100, 1mM EDTA in PBS) at room temperature for 30 min, followed by incubation with primary antibody diluted in blocking solution for 2 hours at room temperature. After washing with PBST (0.1% Tween 20 in PBS) 3 times for 5 minutes each, cells were incubated with Alexa Fluor labeled secondary antibody (Thermo Fisher Scientific) for 45 minutes at room temperature. After washing with PBST 3 times for 5 minutes each, if FISH was performed, cells were dehydrated with ethanol series (70%, 95%, then 100%) and hybridized with TAMRA-OO-(TTAGGG)3 PNA probe (Applied Biosystems) in formamide hybridization solution (70% formamide, 0.5% blocking reagent (Roche), 10 mM Tris-HCl, pH 7.2) at 80°C for 5 min. Cells were allowed to cool at room temperature for 2 hours, then washed 4 time for 10 minutes each with formamide washing solution (70% formamide, 10mM Tris-HCl pH 7.2). Cells were then washed with PBS 3 times for 5 min each, counterstained with DAPI, and coverslips were mounted on slides with anti-fade reagent (Prolong Gold, Invitrogen). Images were captured using a Nikon Eclipse Ti or DeltaVision microscope. The primary antibodies used: mouse Rap1 (1252, rabbit polyclonal), HA (Abcam ab9110, rabbit polyclonal), Flag M2 (Millipore Sigma F1804, mouse monoclonal), Alexa Fluor 488 streptavidin (Thermo Fisher Scientific S11223), and Zscan4 (Millipore Sigma AB4340).

### 2C::tdTomato reporter live-cell imaging

The *2C::tdTomato* reporter was a gift from Samuel Pfaff (Addgene #40281). To establish stable cell lines, mESCs were transfected with the *2C::tdTomato* reporter followed by selection with hygromycin for 7 days to select for cells with stable integrations. Live-cell imaging and analysis was performed using Incucyte (Sartorius). The proportion of *2C::tdTomato* positive cells was determined by normalizing the number of red objects to cell confluency as determined by phase contrast imaging.

### RNA-sequencing

Cell pellets were submitted to Genewiz for RNA extraction, polyA library preparation, and high-throughput sequencing using their pipelines. For bioinformatic analysis of RNA-sequencing (RNA-seq) data, the Seq-N-Slide workflow (source code available at: https://github.com/igordot/sns) was employed. Under this workflow, FASTQ files were trimmed using Trimmomatic (Bolger et al. 2014) to remove adapters and low quality bases and aligned to the mm10 reference genome using RNA-STAR (Dobin et al. 2013). Alignments to other species and common contaminants were removed by fastq_screen (source code available at: https://github.com/StevenWingett/FastQ-Screen). Genes- samples counts matrices were generated using featureCounts (Liao et al. 2019). Differential gene expression analysis was performed by DESeq2 (Love et al. 2014). Differentially expressed genes (DEGs) were determined based on a fold-change (FC) > 1.5 and false discovery rate (FDR) < 0.1. Hierarchically clustered heatmaps were generated using the ComplexHeatmaps package (Gu et al. 2016). Kmeans clustering was performed using Morpheus (https://software.broadinstitute.org/morpheus).

Differential repeat expression analysis was performed as previously described (Ishiuchi et al. 2015). Trimmed RNA-seq reads were aligned to the reference genome using RNA-STAR (Dobin et al. 2013), allowing multimapping of reads. Nonuniquely mapped reads were mapped to the annotation of repeat sequences in the mouse genome using RepeatMasker (http://www.repeatmasker.org/). Differential gene expression analysis was performed by DESeq2 (Love et al. 2014). Differentially expressed repeats were determined based on a FC > 2 and FDR < 0.1. To quantify proportion of DEGs near MERVL MT_Mm sites, BEDtools (Quinlan and Hall 2010) window function was used to count the number of DEGs and total number of genes at increasing distances from MT2_Mm sites as annotated by RepeatMasker

### Chromatin Immunoprecipitation (ChIP)

For Rap1 ChIP, SV40LT-immortalized *Rap1^-/-^* MEFs were transduced with pLPC-Flag- 2xHA-EV, pLPC-Flag-2xHA-Rap1, and pLPC-Flag-2xHA-Rap1-I312R retrovirus produced from transfected Phoenix cells. 10^8^ cells were harvested by trypsinization, washed twice with PBS, and double-fixed with 2mM ethylene glycol bis (succinimidylsuccinate) (EGS, Thermo Fisher Scientific 21565) in PBS for 45 min at room temperature and then with 1% formaldehyde for 20min at room temperature. The reaction was quenched with 125mM glycine and samples were spun 240 x g for 5 min at 4°C. Pellets were washed 2 times with cold PBS, resuspended in 10mL cold lysis buffer 1 (100 mM HEPES-KOH, 140 mM NaCl, 1 mM EDTA, 10% glycerol, 0.5% NP-40, 0.25% Triton X-100, and protease inhibitor cocktail), and incubated rocking for 10 min at 4°C. Samples were spun 950 x g for 2 min at 4°C, pellets were resuspended in 10mL cold lysis buffer 2 (200 mM NaCl, 1 mM EDTA, 0.5 mM EGTA 10 mM Tris-HCl, and protease inhibitor cocktail), and incubated rocking for 10 min at 4°C. Samples were spun 1500 x g for 2 min at 4°C, pellets were resuspended in 3mL cold lysis buffer 3 (1 mM EDTA, 0.5 mM EGTA, 10 mM Tris-HCl at pH 8, 100 mM NaCl, 0.1% Na-Deoxycholate, 0.5% N-lauroyl sarcosine, and protease inhibitor cocktail) and run through a G27 needle 10 times. Samples were sonicated 10 cycles 30sec on/30sec off in a Bioruptor Pico. After sonication, 1/10 volume of 10% Triton X-100 was added and samples were spun 18,400 x g for 10 min at 4°C. Supernatants were precleared with 300uL ChIP-grade Protein G magnetic beads (Cell Signaling Cat. 9006) and then incubated rotating overnight at 4C with 10 µg anti-HA ChIP-grade antibody (Abcam ab9110). The following morning, 30ul ChIP-grade Protein G magnetic beads were added and samples were incubated at 4°C rotating for two hours. Beads were washed a total of 5 times with cold wash buffer (50 mM Hepes at pH 7.6, 1 mM EDTA, 0.7% Na-Deoxycholate, 1% NP-40, 0.5 M LiCl, and protease inhibitor cocktail) and 1 time with PBS. ChIP DNA was eluted in 250 µL elution buffer (0.5% SDS and 100 mM NaHCO3) by incubating at room temperature for 15 min on a roller. A second elution was performed by adding 250 µL elution buffer and incubating 65°C for 20min mixing. Eluates were combined and reverse cross-linked by adding 20 µl 5M NaCl and then incubated at 65C for 4 hours. Samples were then purified by phenol-chloroform-isoamyl alcohol followed by ethanol precipitation and resuspension in water.

For mESC histone marks, 30 x 10^6^ cells were crosslinked in Fixation Buffer (1% formaldehyde, 15mM NaCl, 0.15mM EDTA, 0.075mM EGTA, 10mM HEPES pH 7.6 in DMEM) for 10 min at room temperature, then quenched by addition of 0.125M glycine. Cells were then washed with PBS, harvested by scraping, and centrifuged 2500 x g for 5 min at 4°C. Cell pellets were resuspended in 5mL Lysis Buffer 1 (50mM HEPES pH 7.5, 140mM NaCl, 1mM EDTA, 10% Glycerol, 0.5% NP40, 0.25% Triton-X 100, 10mM Sodium Butyrate, 0.2mM PMSF, and protease/phosphatase inhibitor cocktail), incubated rocking at 4°C for 10 min, then centrifuged 1350 x g for 5 min at 4°C. Cell pellets were then resuspended in 5mL Lysis Buffer 2 (10mM Tris pH 8.0, 200mM NaCl, 1mM EDTA, 0.5mM EGTA, 10mM Sodium Butyrate, 0.2mM PMSF, and protease/phosphatase inhibitor cocktail), incubated rocking at room temperature for 10 min, then centrifuged 1350 x g for 5 min at 4°C. The resulting nuclear pellet was resuspended in 1.2mL Lysis Buffer 3 (10mM Tris pH 7.5, 1mM EDTA, 0.5mM EGTA, 0.5% N-lauroylsarcosine, 10mM Sodium Butyrate, 0.2mM PMSF, and protease/phosphatase inhibitor cocktail) and sonicated for 15 cycles (30 sec on/30 sec off). Sonicated chromatin was spun at high speed for 30 min at 4°C to pellet insoluble debris and supernatant were used for IP. Depending on antibody used, 100 µg of chromatin was aliquoted to a tube, Lysis Buffer 3 was added to bring the volume to 200 µl, and then to 300 µl by addition of 100 µl of Incubation Buffer (3% Triton-X 100, 0.3% Sodium Deoxycholate, 15mM EDTA, 10mM Sodium Butyrate, 0.2mM PMSF, and protease/phosphatase inhibitor cocktail). 8 µg of antibody (anti-acetyl-histone H4, Millipore Sigma 06-598; anti-acetyl-histone H2A.Z, Millipore Sigma Abe1363) was then added and tubes were rotated overnight at 4°C. The next day, Dynabeads Protein A magnetic beads (Thermo Fisher Cat. 1001D) were added and incubated for 2 hours rotating at 4°C. Beads were then washed 5 times with RIPA buffer (50mM HEPES pH 7.5, 0.7% Sodium Deoxycholate, 1mM EDTA, 1% NP40, 500mM Lithium Chloride, 10mM Sodium Butyrate, 0.2mM PMSF, and protease/phosphatase cocktail) and 1 time with 1x TE + 50mM NaCl. Antibody/DNA complexes were then eluted by adding 125 µl Elution Buffer (50mM Tris pH 8.0, 10mM EDTA, 1% SDS) and incubating 20min at 65°C shaking. The eluate was then moved to a new tube and crosslinks were reversed by incubating shaking at 65°C overnight. Then, 4ul of Proteinase K (20mg/mL) was added and tubes were incubated 2 hr at 55°C shaking. ChIP DNA was then purified by QIAquick PCR Purification Kit.

### ChIP dot blot

ChIP DNA was denatured at 95°C for 5 min and dot blotted onto Hybond nylon membranes in 2x SSC. Membranes were treated with 1.5 M NaCl/0.5 N NaOH for 10 min, and with 1 M NaCl/0.5 M Tris-HCl, pH 7.0 for 10 min. DNA was then crosslinked to the membrane using a Stratagene UV crosslinker. End-labeled Telomere probe was prepared by incubating (CCCTAA)_4_ oligo with T4 PNK enzyme and ^32^P-gamma-ATP at 37°C for 45 min. Probe was then filtered through a G-25 column and hybridized to the crosslinked membrane overnight at 55°C. Membranes were washed 3 times in 2x SSC for 5 min, exposed overnight to PhosphorImager screen, and imaged on a Typhoon gel scanner. The quantification of DNA precipitated was performed by measuring integrated intensity using Fiji ImageJ and normalizing to input samples.

### ChIP-sequencing Library Prep, Sequencing, and Analysis

Sequencing libraries were prepared using the NEBNext® Ultra™ II DNA Library Prep Kit for Illumina (NEB Cat. E7645S). For HA-Rap1 ChIP, sequencing was performed on Illumina NextSeq 500 (SE 150 cycles). For histone marks in mESCs, sequencing was performed on Illumina NovaSeq 6000 (PE 100 cycles).

For bioinformatic analysis of ChIP-sequencing (ChIP-seq) data, the Seq-N-Slide workflow (source code available at: https://github.com/igordot/sns) was employed. Under this workflow, reads were aligned to the mm10 reference genome using Bowtie2 (Langmead and Salzberg 2012), duplicate reads were removed using Sambamba (source code available at: https://github.com/biod/sambamba), and genome browser tracks representing ChIP-seq profiles were generated using IGV (Robinson et al. 2011). MACS2 (Zhang et al. 2008) was used for peak calling. Deeptools (Ramirez et al. 2016) was used to generate heatmaps and density plots.

For telomere binding analysis using high-throughput sequencing, FASTQ files were processed using grep to count the number of reads containing *≥*3 consecutive telomere repeats (TTAGGG/CCCGAA) and normalized to the total number of reads.

### qRT-PCR

RNA was purified from cells using the NucleoSpin RNA Mini Kit (Macherey-Nagel. 740955.50). 1ug of RNA was reverse transcribed using the iScript gDNA Clear cDNA Synthesis Kit (BioRad 1725034) and cDNA libraries were diluted 1:5. RT-PCR was performed using ssoAdvanced SYBR Green Supermix (Cat. 1725270) on a Roche LightCycler 480 II. qPCR primers can be found in Supplementary Table S6.

### Subcellular Fractionation

1 x 10^7^ cells were resuspended in 200 µL Buffer A (10mM HEPES pH 7.9, 10mM KCl, 1.5mM MgCl, 0.34M Sucrose, 10% Glycerol, 1mM DTT, 1mM PMSF, and protease inhibitor cocktail). After addition of 2uL of 10% Triton-X 100, resuspension was incubated 8 min on ice. Samples were centrifuged 1300 x g for 5 min at 4°C. Supernatants (S1) were clarified at 20,000 x g for 5 min at 4°C to yield cytoplasmic fraction (S2). The pellet washed once with Buffer A and then lysed in 100 µL Buffer B (3mM EDTA, 0.2mM EGTA,

1mM DTT, 1mM PMSF, and protease inhibitor cocktail) for 30 min at 4°C. Samples were centrifuged 1700 x g for 5 min at 4°C and the supernatant was kept as the nucleoplasmid (S3) fraction and pellets were the chromatin bound fraction (P3). Chromatin was washed one time with Buffer B and then resuspended in Laemmli buffer, sonicated, and boiled for 10min at 70°C.

### Nucleosome reconstitution

The Widom 601 nucleosome positioning sequence was obtained from a plasmid containing 8 copies of the fragment, each flanked by the EcoRV restriction site (Armache et al. 2011). Wild-type *Xenopus* histone was expressed individually in *E. coli* cells, extracted from inclusion bodies, and purified by size exclusion and anion chromatography (Dyer et al. 2004b). Purified histones were lyophilized (Sentry, VirTis) and stored at -80C for further use. Nucleosomes reconstitutions were performed as described (Dyer et al. 2004a; Armache et al. 2011). Briefly, recombinant histone octamers were assembled by mixing equimolar amounts of each of the four histones and dialyzing against refolding buffer (10 mM Tris, pH 7.5, 2.0 M NaCl, 1 mM EDTA, 5 mM BME). Assembled octamers were purified by size exclusion chromatography (Superdex 200, GE healthcare) in refolding buffer. Then, histone octamers and purified Widom 601 DNA were combined and dialyzed overnight with salt gradient dialysis (Rapid Pump, Gilson). Finally, assembled nucleosomes were purified through ion-exchange chromatography (Resource Q, GE Healthcare). Purified nucleosomes were dialyzed into TCS buffer (20 mM Tris-HCl pH 7.5, 1 mM EDTA, 1 mM DTT), concentrated, and stored at 4°C.

### Proximity-based biotinylation (BioID) and streptavidin pull-down

SV40LT-immortalized *Rap1^-/-^* MEFs were transduced with pLPC-Flag-BirA*-13xGGGGS- Myc-EV, pLPC-Flag-BirA*-13xGGGGS-Myc-Rap1, and pLPC-Flag-BirA*-13xGGGGS- Myc-Rap1-I312R retrovirus produced from transfected Phoenix cells. Single-cell clones were isolated, and independent clonal cell lines were screened by IF to identify those that expressed at high levels. Cells grown in 15-cm^2^ dishes to be 90% confluency at time of harvest were treated with biotin (50 µM) for 20 hours followed by biotin free media for 1hr. For each condition, 10 x 10^6^ cells were harvested by trypsinization, lysed for 1hr rocking at 4°C in NP40 lysis buffer (10mM Tris pH 7.4, 10mM NaCl, 3mM MgCl_2_, 1mM PMSF, and protease inhibitor cocktail), and then centrifuged 3300 x g 10min at 4°C to pellet nuclei. Nuclei were resuspended in 500ul SDS Lysis Buffer (50mM Tris pH 8.0, 1% SDS, 10mM EDTA, 1mM PMSF, and protease inhibitor cocktail), incubated on ice for 10min, followed by boiling for 5 min at 95°C. Samples were then sonicated for 10 cycles (30 sec on/30 sec off) and clarified by centrifuging 15,000 x g for 10 min at 4°C. Lysates were clarified by centrifuging 15,000 x g for 10 min at 4°C. 50uL MyOne Streptavidin Beads (Thermo Fisher 65001) were then added and incubated rotating at 4°C overnight. Beads were then washed 2 times with RIPA buffer, 2 times with 2M Urea in 10mM Tris pH 8.0, 2 more times with RIPA, and then 2 times with Buffer 4 (50mM Tris pH7.4, 50mM NaCl). Proteins were then eluted from magnetic beads into Laemmli Buffer + 2mM biotin by boiling 5min at 95°C. Eluates were used for Western blot, Silver stain (Thermo Fisher Scientific 24612), and mass spectrometry analysis as previously described (Pinzaru et al. 2020).

### Coimmunoprecipitation (CoIP)

HEK293T cells were cotransfected with the indicated plasmid constructs using polythelenimine (PEI). 48hr post-transfection, cells were harvested in cold PBS, washed once with PBS, and lysed for 15min on ice in 500 µL Lysis Buffer (50mM Tris-HCl pH 7.4, 150mM NaCl, 1% Triton X-100, 0.05% SDS, 1mM EDTA, 1mM DTT, 1mM PMSF, and protease inhibitor cocktail). Salt boost was performed by adding 25 µL 5M NaCl and incubating on ice additional 5min. Then, 500 µL cold H_2_O was added and lysates were spun at max speed for 10min to pellet debris. Antibody for epitope to be precipitated (anti-Flag, Millipore Sigma F1804; anti-HA, BioLegend 901533; anti-c-Myc Millipore Sigma M4439) was added to supernatants and incubated for 3 hours rotating at 4°C. Protein G Sepharose (Millipore Sigma Cat. GE17-0618-01) preblocked in 5% BSA was added and incubated for 1 additional hour rotating at 4°C. Beads were washed 4 times in Wash Buffer (50mM Tris-HCl pH 7.4, 150mM NaCl, 1% Triton X-100, 0.1% SDS) and proteins were eluted in Laemmli buffer by boiling at 95°C for 5min for Western blot or with 3xFlag peptide (150 µg/ml, Sigma) for 1 hr at 4°C for HAT assay. For CoIP of Rap1 with K48-Ub, the protocol was the same as above except no salt boost was performed, TNE (10 mM Tris pH 7.8, 1% NP-40, 0.15 M NaCl, 1 mM EDTA, and protease inhibitor cocktail) was used as lysis and wash buffer, and 4 mM N-ethylmaleimide (Sigma Cat. E3876) and 4 mM 1,10-phenanthroline (Sigma Cat. 131377) were included in the buffers. For DNase sensitivity assays, lysis buffer did not include EDTA or DTT. Lysates were supplemented with MgCl2 (2.5 mM) and incubated with DNase (10 µg/mL) for 1hr at RT. DNase was then inactivated by addition of EDTA and DTT before addition of antibody for IP.

### Affinity purification of Rap1 protein

Recombinant Rap1 with N-terminal His_6_-tags was expressed from the pTriEx4 vector in BL21(DE3) RIPL bacterial cells as described elsewhere (Vizlin-Hodzic et al. 2009; Nora et al. 2010). After 3-hour long induction of protein expression with 0.5mM isopropyl *β*-D- thiogalactoside, cells were harvested and resuspended in lysis buffer (50 mM sodium phosphate, 300 mM NaCl, 10 mM imidazole, 10% glycerol, and 0.5% Tween 20, pH 8.0) containing protease inhibitor cocktail (Roche). Sonicated cell extracts were cleared by centrifugation and subsequently filtered (0.45 µm SterivexTM filter, Millipore). The supernatant containing Rap1 was further purified by Immobilized Metal ion Affinity Chromatography using TALON Metal Affinity Resin (Clontech) containing Co2+ cations as described (Yanez et al. 2005). Rap1 containing fractions were eluted in 200 mM imidazole. For His_6_-tag removal, samples were digested by thrombin protease and purified using gel filtration chromatography to remove cleaved tags and residual protease. The fractions containing pure protein were dialyzed into buffer composed of 50 mM sodium phosphate with 50 mM NaCl (pH 7.0) and, subsequently, proteins were concentrated by ultrafiltration (Amicon 30K, Millipore). The concentration of purified proteins was determined using the Bradford assay and purity was verified by electrophoresis in sodium dodecyl sulfate (SDS)-polyacrylamide gel which was subsequently stained using Bio-Safe Coomassie G 250 (Bio-Rad).

### Electrophoretic mobility shift assay (EMSA)

Increasing amounts of 6xHis-tagged Rap1 (0 to 15 µM) or tag-free Rap1 proteins (0 to 15 µM) were incubated with 100 nM DNA (74bp long telomeric or 601 sequence) or nucleosome substrate in EMSA buffer (10 mM Tris pH 7.5, 100 mM NaCl, 2.5% Glycerol and 1 mM DTT) at room temperature for 30 min. The binding reactions were resolved on native polyacrylamide gels (6% PAGE, 0.25 x TBE). Runs were performed in the cold room (4°C) at 170 V for 1.5 hour, gels were stained with ethidium bromide and visualized on a Typhoon Trio+ scanner (Molecular Dynamics). Purified nucleosome substrates were a generous gift from Dr. Karim-Jean Armache.

### Histone acetyltransferase (HAT) assays

Core histones and short oligonucleosome substrates for HAT assays were purified as previously described (Cote et al. 1995). Indicated substrates were incubated in a 15 µl reaction containing 50mM Tris-HCl pH8.0, 10% glycerol, 1mM EDTA, 1mM DTT, 1mM PMSF, 10mM sodium butyrate and 0.125 μCi of [3H]-labeled Acetyl-CoA (Perkin Elmer). Samples were then spotted on phosphocellulose paper (St Vincent’s Institute) and counts per minute (CPM) were quantified using Beckman Coulter LS6500 Multipurpose Scintillation Counter.

### Tandem affinity purification (TAP) of native TIP60/p400 complex for HAT assays

Native TIP60/p400 complex was affinity purified as previously described (Dalvai et al. 2015). Briefly, nuclear extracts (Abmayr et al. 2006) were prepared from 1 x 10^9^ – 3 x 10^9^ K562 cells expressing EPC1-3xFLAG-2xStrep from the *AAVS1* locus (empty vector for Mock preparations). Tween-20 was added to 0.1%, and extracts were centrifuged 17,000 rpm in Beckman JA-20 rotor for 1h at 4C. After preclearing with 300ul Sepharose CL-6B (Sigma), extracts were incubated with 250uL anti-Flag M2 affinity resin (Sigma) for 2 hours at 4C. Beads were then washed in Poly-Prep columns (Bio-Rad) with 40 column volumes of Buffer 1 (20 mM HEPES-KOH pH 7.9, 10% glycerol, 300 mM KCl, 0.1% Tween 20, 1 mM DTT, Halt protease and phosphatase inhibitor cocktail without EDTA (Pierce)), followed by 40 column volumes Buffer 2 (20 mM HEPES-KOH pH 7.9, 10% glycerol, 150 mM KCl, 0.1% Tween 20, 1 mM DTT, Halt protease and phosphatase inhibitor cocktail without EDTA (Pierce)). Protein complexes were eluted with 3xFlag peptide (200ug/mL, Sigma) in 5 column volumes Buffer 2 for 1 hour at 4C. The eluted fraction was then mixed with 125ul Strep-Tactin Sepharose (IBA) affinity matrix for 1 hr at 4°C. Beads were then washed again in in Poly-Prep columns (Bio-Rad) with 40 column volumes of Buffer 2. Complexes were then eluted in two fractions with 2.5mM D-biotin in 4 column volumes Buffer 2 and flash frozen in liquid nitrogen.

### siRNA Transfection

For mESCs, 75,000 cells were reverse transfected into gelatin-coated 24-well plates using Lipofectamine RNAiMAX (Thermo Fisher Scientific Cat. 13778100) according to the manufacturer’s instructions and a final concentration of 30nM for each siRNA. 48hr post-transfection, RNA was purified from cells using the NucleoSpin RNA Mini Kit (Macherey-Nagel Cat. 740955.50). A list of siRNAs used in this study can be found in Supplementary Table S7.

### RAP1-EPC1 interaction *in vitro*

1ug of soluble His-Rap1 was pre-cleared on Glutathione sepharose 4B beads (GE Healthcare) for 1h at 4C, then incubated with either 3µg (measured by Bradford protein assay) of GST-tagged EPC1 (aa 1-581) or GST alone immobilized on Glutathione sepharose beads for 2h30min at 4C in 200uL Pulldown Buffer [200mM NaCl, 25mM Hepes pH 7.5, 10 % glycerol, 100 µg / mL BSA, 1 mM PMSF, 0.5 mM DTT, 0.1 % Tween- 20, 2 µg / mL leupeptin, 2 µg / mL pepstatin, 5 µg / mL aprotinin]. Beads were washed twice with Pulldown Buffer and ran on SDS-PAGE followed by western blotting (anti-His Clontech 631212).

